# Spatial single-cell profiling of intracellular metabolomes *in situ*

**DOI:** 10.1101/510222

**Authors:** Luca Rappez, Mira Stadler, Sergio Triana, Prasad Phapale, Mathias Heikenwalder, Theodore Alexandrov

**Affiliations:** Structural and Computational Biology Unit, European Molecular Biology Laboratory (EMBL), Heidelberg, 69117 Germany; Collaboration for joint PhD degree between EMBL and Heidelberg University, Faculty of Biosciences, Germany; Institute of Chronic Inflammation and Cancer, Deutsches Krebs-Forschungszentrum (DKFZ), 69120 Heidelberg, Germany; Metabolomics Core Facility, EMBL, Heidelberg, 69117 Germany; Skaggs School of Pharmacy and Pharmaceutical Sciences, University of California San Diego, CA 92093, La Jolla, USA

**Keywords:** Spatial single-cell metabolomics, imaging mass spectrometry, microscopy, heterogeneity, metabolic intermixing, macrovesicular steatosis, lipid droplets, cell-cell contact, SpaceM

## Abstract

The recently unveiled extent of cellular heterogeneity demands for single-cell investigations of intracellular metabolomes to reveal their roles in intracellular processes, molecular microenvironment and cell-cell interactions. To address this, we developed SpaceM, a method for *in situ* spatial single-cell metabolomics of cell monolayers which detects >100 metabolites in >10000 individual cells together with fluorescence and morpho-spatial cellular features. We discovered that the intracellular metabolomes of co-cultured human HeLa cells and mouse NIH3T3 fibroblasts predict the cell type with 90.4% accuracy and revealed a short-distance metabolic intermixing between HeLa and NIH3T3. We characterized lipid classes composing lipid droplets in steatotic differentiated human hepatocytes, and discovered a preferential accumulation of long-chain phospholipids, a co-regulation of oleic and linoleic acids, and an association of phosphatidylinositol monophosphate with high cell-cell contact. SpaceM provides single-cell metabolic, phenotypic, and spatial information and enables spatio-molecular investigations of intracellular metabolomes in a variety of cellular models.

## Introduction

Multicellular organisms contain a multitude of cells of distinct and diverse functions, morphologies, and molecular compositions. Each single cell has a unique intracellular metabolome, a dynamic repertoire of metabolites and lipids involved in virtually all cellular processes. Aside with metabolites and lipids serving as building blocks and energy sources within the cell, recent discoveries unveiled their roles in signaling (Wellen and Thompson, 2012), epigenome regulation (Sharma and Rando, 2017), immunity (Buck et al., 2017), inflammation (Murphy and O’Neill, 2018), host-microbe interactions (Sharon et al., 2014), and cancer (Pavlova and Thompson, 2016). At the same time, the progress of single-cell technologies revealed the extent and biological functions of cellular heterogeneity (Altschuler and Wu, 2010) within tissues, organs (Marioni and Arendt, 2017), tumors (Patel et al., 2014), and even among monoclonal cells in culture (Lee et al., 2014; Pelkmans, 2012; Russell et al., 2018). The discovered critical roles of metabolism and the growing awareness of a hidden world beneath population averages created an urgent need to investigate intracellular metabolism at the single-cell level (Rubakhin et al., 2013; Zenobi, 2013). In the recent years, the sensitivity of mass spectrometry-based metabolite detection has improved substantially opening novel avenues to metabolomics of either single cells or small groups of cells (Do et al., 2017; Guillaume-Gentil et al., 2017; Ibáñez et al., 2013; Merrill et al., 2017) and even at a subcellular level (Passarelli et al., 2017). However, despite these methods successfully demonstrated detection of metabolites in individual cells, analytical and computational challenges precluded studies of spatio-molecular organization and cellular heterogeneity, and prevented discovering the links between intracellular metabolomes, cellular phenotypes and spatial organisation of cells.

To bridge this gap, we designed SpaceM, a method for spatial single-cell metabolomics of cell monolayers that integrates MALDI-imaging mass spectrometry with bright-field and fluorescence microscopy. Integration with microscopy enables associating metabolites with fluorescence and morphological cell properties (fluorescent reporter intensity, area, compactness, shape) as well as with spatial features quantifying multi-cellular organization. The integration was enabled by a method for precise detection of parts of cells sampled by MALDI laser with the help of sequential microscopy, novel image analysis, and a novel cell-ablation marks normalisation strategy. Using the False Discovery Rate-controlled metabolite annotation, and novel methods for unbiased selection of intracellular metabolites and for filtering out poor quality cells allowed us to perform high-throughput analyses with >100 metabolites detected in >10000 individual cells, with a high reproducibility between replicates. We validated SpaceM by investigating metabolomes of co-cultured HeLa and mouse fibroblasts cells as well as of differentiated human steatotic hepatocytes stimulated with pro-inflammatory factors that provided rich metabolic, phenotypic, and spatial information.

## Results

### The SpaceM method

SpaceM relies on using Matrix Assisted Laser Desorption Ionization (MALDI)-imaging mass spectrometry, a spatially-resolved mass spectrometry technology for detection of a wide range of molecules (Baker et al., 2017). MALDI-imaging is increasingly used for spatial metabolomics (Palmer et al., 2016) and was demonstrated to achieve the femtomolar-levels sensitivity (Soltwisch et al., 2015). This, together with soft ionisation preventing excessive in-source molecular fragmentation makes it a perfect choice for single-cell metabolomics as demonstrated by others (Do et al., 2017; Ibáñez et al., 2013). The experimental part of SpaceM combines MALDI-imaging with microscopy as well as with collecting supporting information to integrate these two sources of data (Figure 1; for a detailed workflow see Figure S1). The cells for SpaceM are cultured on a labtek chamber glass slide in a monolayer, with the cell confluence sufficient to allow cells to interact between each other but at the same time preventing the growth of cells on top of each other. After washing, cells are fixed to halt enzymatic activity, stained with a fluorescent dye with the staining protocol compatible with metabolomics, and dried in a desiccator following regular cell preparation protocols. SpaceM requires the Hoechst (or any similar) staining for nuclei detection. For investigation of the steatotic hepatocytes, we also used the lipophilic LD540 staining to detect lipid droplets (Spandl et al., 2009). Then, bright-field and fluorescence microscopy images of cells are collected with the following two aims in mind.

**Figure 1.**
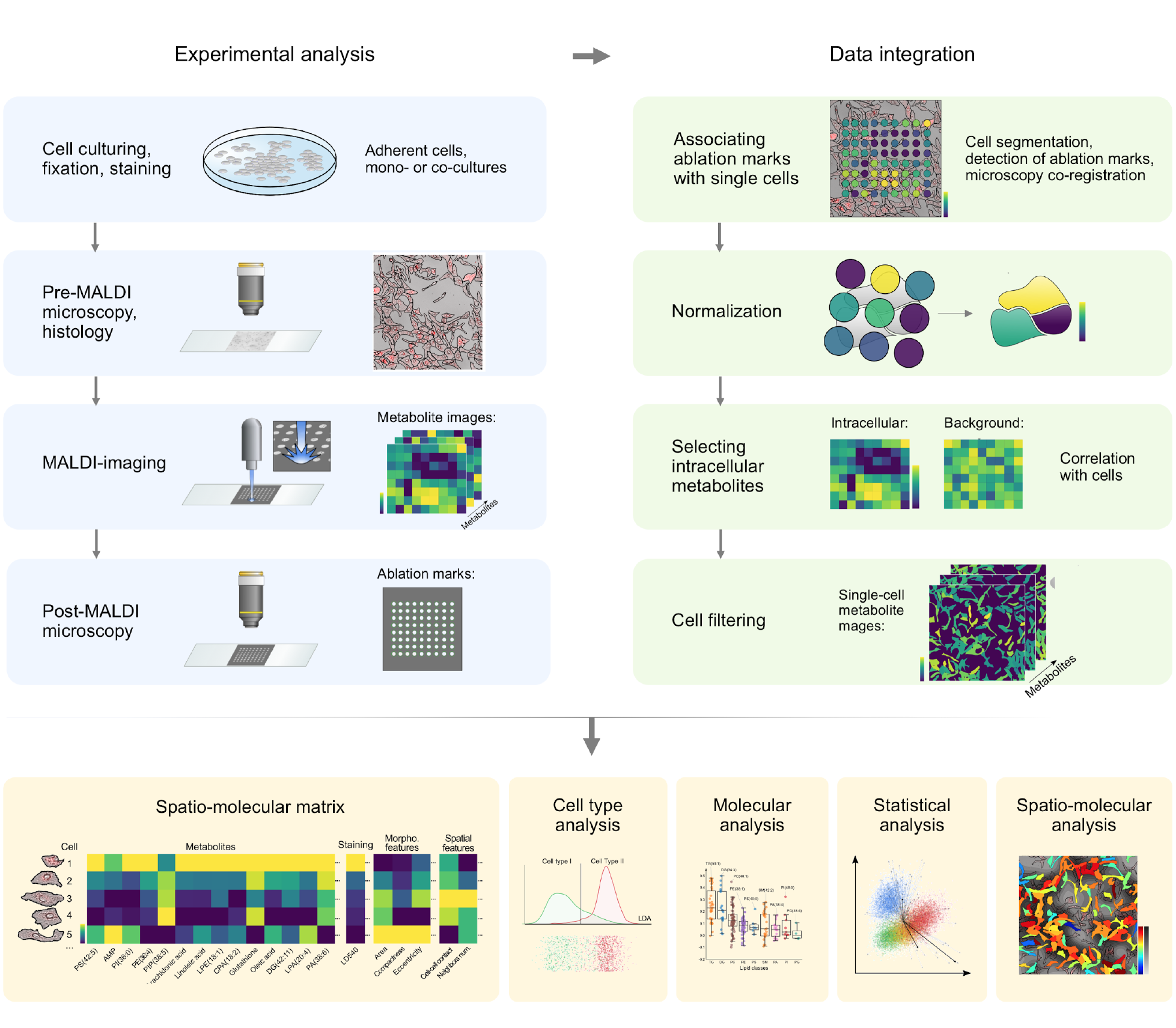
SpaceM method for spatial single-cell metabolomics of cell monolayers by integrative microscopy and MALDI imaging mass spectrometry. The experimental part of the workflow includes cell culturing, pre-MALDI and post-MALDI microscopy and histology, and MALDI imaging mass spectrometry. The data integration part includes associating of MALDI laser ablation marks with individual cells, strategies for normalization, selecting intracellular metabolites, and cell filtering (see Figure S1 for a detailed workflow). SpaceM outputs a single-cell spatio-molecular matrix providing rich information for a variety of analyses, in particular to characterise cell types, associate single-cell metabolomes with a fluorescent phenotype,interrogate changes of single-cell metabolomes upon perturbations, and discover spatio-molecular associations.

First, the cell segmentation of the microscopy images provides cell outlines and enables cell localization. Second, microscopy provides rich phenotypic information about single-cell fluorescence, immunochemistry and spatio-morphological properties of the cells. In the next experimental step, MALDI-imaging is applied to the dried cells to collect mass spectra across cells and extracellular areas. MALDI-imaging procedure starts with application of an ionisation-enhancing matrix. Similar to MALDI-imaging of tissues, we used a robotic sprayer for enhanced extraction, high spatial resolution, and high reproducibility. MALDI-imaging generates big datasets with millions of mass-to-charge channels. For finding metabolic signals in this data, we exploited the False Discovery Rate-controlled metabolite annotation implemented as the METASPACE cloud software (http://metaspace2020.eu) (Palmer et al., 2017). METASPACE is an essential step as it reduces millions of mass-to-charge (m/z)-values to ~100 metabolite annotations, filters out signals representing matrix and contaminants, ensures quality control and represents metabolite images a user-friendly way. In the last experimental step, we performed post-MALDI microscopy to determine which cells were sampled by the MALDI-imaging laser and to associate MALDI-imaging spectra with the cells. Next, we performed data integration with the first step associating ablation marks with individual cells. We detected MALDI laser ablation marks in post-MALDI microscopy images using a customized 2D Fourier Transformation image analysis method that exploits similarities between ablation marks and the regularity in spacings between them. Then, we obtained positions of the MALDI-imaging ablation marks within the cell areas by co-registering pre-MALDI microscopy images (containing cell outlines) with post-MALDI microscopy images (containing ablation marks outlines) (Figure S2). For a majority of cells, a cell was sampled with just one ablation mark. In our benchmarking experiment with HeLa cells and NIH3T3 fibroblast (described later), 72.25%, 23.8%, 2.8%, 0.9% cells were sampled with 1, 2, 3, 4 ablation marks, respectively. To integrate metabolic profiles from several ablation marks co-sampling the same cell as well as to reduce the confusion of co-sampled cells, we developed a cell-ablation marks normalization strategy (Figure 2). The normalization provides the metabolite intensity normalized by the area that is a natural readout for metabolite concentration. Next, we developed a strategy to distinguish intracellular from extracellular metabolites by requiring an intracellular metabolite intensities to be highly correlated with the cell spatial distribution. Finally, similar to cell filtration strategies in single-cell RNA-seq (Ilicic et al., 2016), we filter out poor-quality cells with low numbers of metabolite annotations with the cutoff determined as the 5%-percentile of the numbers of annotations for all cells in an experiment.

**Figure 2.**
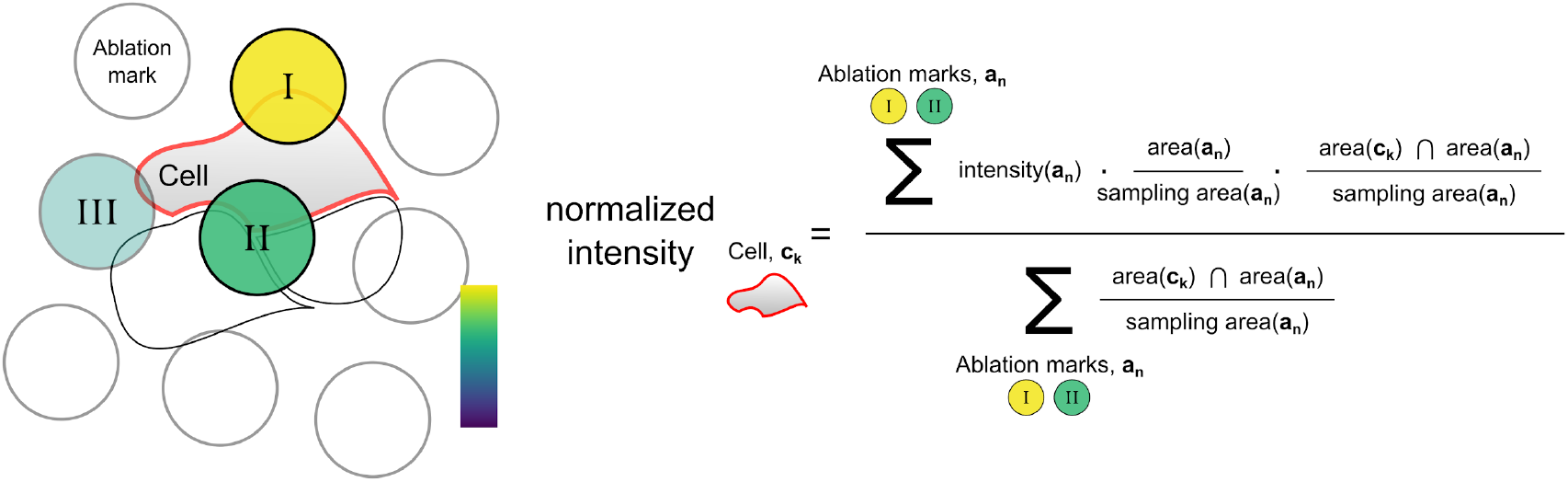
Normalization strategy for assigning metabolic intensities to individual cells. The intensity assigned to a cell for a given metabolite is calculated as a weighted mean of the metabolite intensities from the ablation marks sampling that cell. To increase the contribution of ablation marks which mainly sample the cell of interest, the weight of each ablation mark is proportional to the overlap of the ablation mark and the cell. To reduce the contribution of ablation marks which mainly sample extracellular areas, the weight of each ablation mark is reversely proportional to its extracellular sampling area. Ablation marks sampling predominantly extracellular areas are filtered out (as the illustrated here ablation mark III).area(a_n_) stands for the area of ablation mark an; sampling area(an) stands for the intracellular area of ablation mark an; area(ck) stands for the area of cell c_k_; all areas are computed in microscopy pixels.

Ultimately, SpaceM provides a single-cell spatio-molecular matrix that, for each cell, comprises a multiplex readout of the cell metabolic profile, fluorescence intensity, morphological features, and spatial features (Figure 1). This information enables statistical analysis, phenotype-metabolome correlation, and/or spatio-molecular interrogation of single cells in the genuine spatial context.

### SpaceM predicts cell types with single-cell resolution

For validation of the method, we evaluated whether SpaceM can predict the cell type of spatially-heterogeneous co-cultured human HeLa cells and mouse NIH3T3 fibroblasts (Figure 3). We considered six samples (two replicates of co-cultures, and two replicates of control monocultures for each cell type). HeLa and NIH3T3 cells constitutively expressed H2B-mCherry and GFP, respectively, making them easily discernible by fluorescence microscopy. Automated assignment of the cell type was done using a linear separating boundary between mCherry and GFP fluorescence intensities (Figure 3A). Overall, metabolic profiles of 88 metabolites with an FDR<10% in at least one sample were obtained for 1624 cells in co-cultures (958 HeLa and 666 NIH3T3 cells) and 2197 cells in monocultures (1603 HeLa and 594 NIH3T3 cells). Support Vector Machine classification of the single-cell metabolic profiles with the 10-fold cross-validation for unbiased choice of the Gaussian kernel gamma predicted the cell type with 90.4% accuracy for the co-cultured cells. To evaluate the richness of the detected metabolic profiles, we used only a random subset of all 88 metabolites and were able to consistently predict the cell type in co-cultures with 87.6% accuracy even when using only a half of all detected metabolites (Figure 3B). This indicates the richness of the detected intracellular metabolomes and its biological relevance in the context of cell type classification. Moreover, by applying the statistical t-test we have identified the molecular markers of each cell type in co-cultures, with the phosphatidylinositols PI(34:1), PI(34:2) found to be the most statistically significant and exhibiting the highest fold change in HeLa cells.

**Figure 3.**
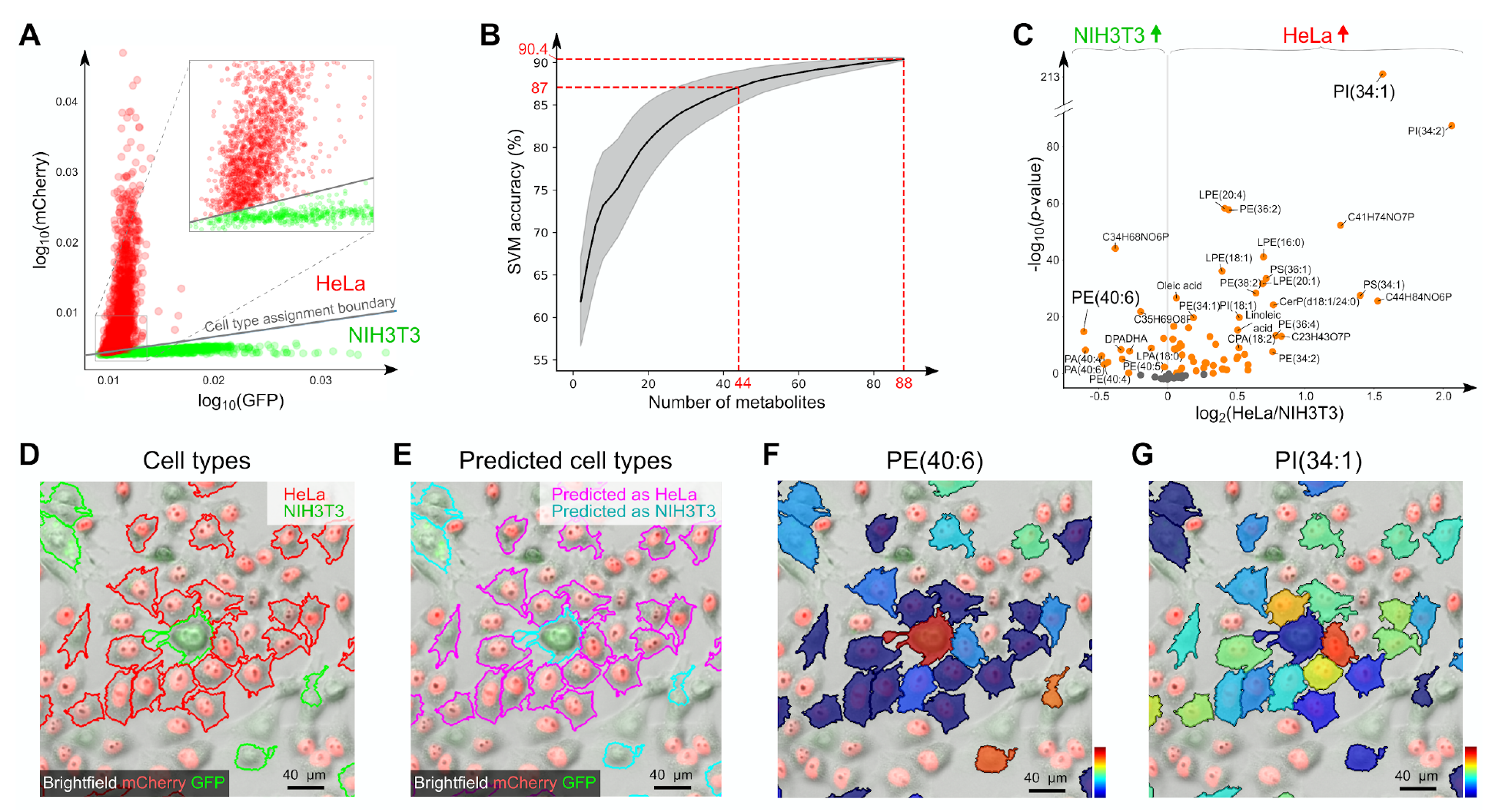
SpaceM can predict the cell type of co-cultured HeLa cells and NIH3T3 fibroblasts based on their intracellular metabolomes, with single-cell resolution. **A:** Automated assignment of the cell type for co-cultured HeLa and NIH3T3 cells (n=3821) was done using a linear separating boundary between mCherry and GFP fluorescence intensities. **B:** Using single-cell metabolic profiles, we could predict the cell type with 90.4% accuracy; even when using only half of the detected metabolites, we could predict the cell type with 87.6% accuracy on average indicating the richness and the relevance of the detected metabolomes; Support Vector Machine with 10-fold cross validation was used for classification; the plot shows the median accuracy for 1000 random repetitions when subsampling the metabolites, with the grey area showing the confidence intervals of ± one standard deviation. **C:** Volcano plot (log2 of the fold change HeLa/NIH3T3 vs. −log10 of the t-test p-value) showing differential properties 88 detected metabolites and lipids. **D:** Area of the co-cultured cells showing an NIH3T3 cell surrounded by HeLa cells; the sampled cells are outlined according to their assigned type. **E:** SpaceM demonstrates single-cell resolution allowing correct prediction the cell type of the NIH3T3 cell surrounded by HeLa cells. **F-G:** Single-cell metabolite images for two markers highlighted in Figure 2E: phosphoethanolamine PE(40:6) and phosphatidylinositol PI(34:1) validated using LC-MS/MS (Data S1); despite the single-cell metabolite images exhibiting visually-noticeable heterogeneity, the full metabolic profile let predict the cell type with a high accuracy indicating the richness of the detected metabolomes and their relevance for each cell type.

In order to assess whether the metabolic profiles can predict the cell type with a single-cell resolution, we evaluated a case of a fibroblast surrounded by HeLa cells (Figure 3D). As shown in Figure 3E, its cell type was predicted correctly. The phosphoethanolamine PE(40:6), the fibroblast marker for NIH3T3 with the highest fold change (Figure 3F) is exhibiting a higher intensity in the fibroblast compared to the surrounding HeLa cells. Interestingly, the visualization of the intensities of the most significant HeLa marker PI(34:1) shows high cell-cell heterogeneity among the HeLa cells (Figure 3G), confirming that it is a combination of several metabolites that enables prediction of the HeLa cell type.

### Intermixing of metabolomes of co-cultured HeLa and NIH3T3

Intriguingly, the detected intracellular metabolomes of co-cultured cells were different from metabolomes of their monocultured counterparts (Figure 4). The co-culturing was optimized to achieve high spatial heterogeneity so that cells of one type have neighbors of another type (Figure 4A-B). We observed that upon co-culturing, the single-cell metabolomes of one cell type become more similar to the single-cell metabolomes of another cell type as compared to their monocultured counterparts (Figure 4C). The values of the metabolic discriminant do not overlap between the monocultured HeLa and NIH3T3 (Figure 4C, with the 90% confidence interval shown monocultured cells of each type). However, for the co-cultured cells, 39.2% of NIH3T3 cells (and 48.2% of HeLa cells) exhibit the values of the metabolic discriminant in the mixing region between the cell types or even take the values within the confidence interval of the other type. This observation of metabolic intermixing between co-cultured cells was further supported by results from predicting the cell type based on single-cell intracellular metabolomes. Whereas we could predict the cell type of co-cultured cells with 90.4% accuracy, for the monocultured cells we could predict their cell type with a higher accuracy of 96.6%.

**Figure 4.**
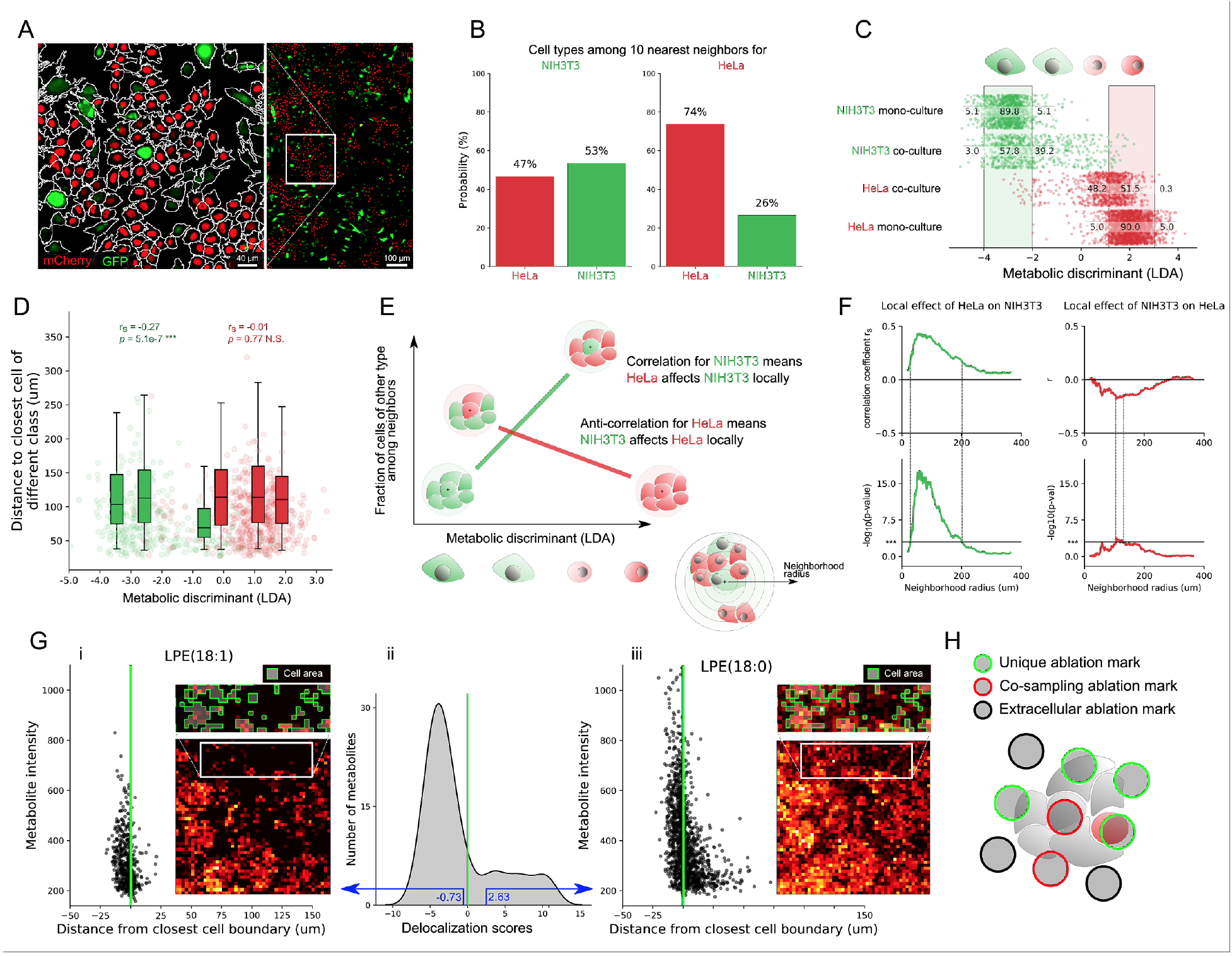
Metabolic intermixing of co-cultured HeLa and NIH3T3 fibroblasts. **A:** Illustration of the spatial heterogeneity of co-cultured HeLa (H1B-mCherry, red) and NIH3T3 (GFP, green). **B:** Quantification of the spatial heterogeneity showing that, for each cell type, individual cells are surrounded by cells of other type (on average, an NIH3T3 cell has 47% HeLa among 10 closest neighbors, a HeLa cell has 26% NIH3T3). **C:** Less pronounced difference between HeLa and NIH3T3 cells in co-cultures compared to their monocultured counterparts suggests metabolic intermixing between the cell types, here visualized with the values of the Linear Discriminant Analysis (LDA) discriminant of the single-cell metabolic profiles; the cells are colored and plotted against its cell type and whether they were cultured in co- or mono-culture; the cell type for co-cultured cells was determined using their fluorescence; the vertical bars indicate the 90%-confidence intervals for each monocultured cell type. **D:** The metabolomes of NIH3T3 cells are affected most when they have HeLa cells in close vicinity with distance between cell centers of 50-100 µm, whereas the average diameter of a HeLa cell equal to 6.7 (+/- 1.7) pm and of NIH3T3 equal to 8 (+/- 2.47) µm. **E:** Illustration explaining the calculation of the distance of the metabolic intermixing effect, per cell type: For each cell, we consider a neighborhood of particular radius, and calculate a fraction of cells of the other type in this neighborhood. The radius for which this property is most correlated (for NIH3T3 cells) or anti-correlated (for HeLa cells) with the metabolic discriminant (LDA) corresponds to the distance of the intermixing effect. **F:** NIH3T3 cells are most affected by HeLa at the distance of 58 µm; HeLa cells are mainly affected by NIH3T3 cells at the distance of 120 µm, with the correlation coefficient and p-values indicating the smaller extend of this effect compared to NIH3T3. **G:** Evaluation of the metabolite delocalization confirmed that the metabolic intermixing cannot be explained by the metabolite delocalization between neighboring cells. The scatterplots for ablation marks for lysophosphoethanolamines LPE(18:1) (i) and LPE(18:0) (iii) show how their metabolite intensities depend on the distance to the nearest cell boundary (negative for intracellular ablation marks, positive for extracellular ablation marks). LPE(18:0) exemplifies a delocalized metabolite with the high metabolite intensities observed at the extracellular ablation marks at the distances of <50 µm from cells. LPE(18:1) exemplifies a well-localized metabolite detected predominantly in the intracellular ablation marks only. ii) a histogram of the delocalization scores for all metabolites showing that most of the metabolites are well-localized; moreover, when considering only well-localized metabolites, SpaceM predicts the cell type with the classification accuracy 88.9% suggesting that despite a minority of metabolites (23 out of 88) having positive delocalization scores, the metabolic intermixing between the cell types is not explained by the delocalization. **H:** We confirmed that the metabolic intermixing is not due to the co-sampling of cells (when an ablation mark samples more than one cell e.g. as illustrated with red outlines), since even after considering only well-localized metabolites and excluding co-sampled SpaceM could predict the cell type with 90.7% accuracy (cf. 90.4% for all metabolites and cells). *** denotes p-values<0.001; N.S. stands for non-significant.

Furthermore, we discovered that the metabolic intermixing between cells of two types happens locally and can be considered a short-distance effect. The extent of the metabolic intermixing for NIH3T3 cells depends on the presence of HeLa in their close vicinity (Figure 4D). We estimated that the metabolic intermixing is the strongest at the distance of 58 µm for NIH3T3 and 107 µm for HeLa, with NIH3T3 affected most (rs=0.43, −log10(*p*-value)=17.5 compared to rs=-0.18, − log10(*p*-value)=3.8). We evaluated whether the observed metabolic intermixing can be observed either due to metabolite delocalization during sample preparation or due to co-ablation of cells. Sample preparation is known to be a key to achieve high spatial resolution MALDI-imaging. Particularly critical is the application of the MALDI matrix, since it can cause metabolite delocalization either during matrix crystallization as crystals can contain analytes from the whole crystal footprint, or due to metabolite leakage while spraying the matrix solution. We have developed a strategy to quantify metabolite delocalization (Figure 4G) that showed that most of metabolites (65 of 88) are well-localized. The delocalized metabolites such as lysophosphoethanolamine LPE(18:0) (see Figure G, cf. well-localized LPE(18:1)) showed only minor levels of delocalization with median distance outside of cells <5 µm. Importantly, even when using only well-localized metabolites, SpaceM could predict the cell type with 88.9% accuracy. The fact that the cell type prediction accuracy did not increase after considering localized metabolites only suggests that delocalization does not explain the observed intermixing between the cell types. Next, we evaluated whether co-ablation of cells can be a reason for the observed metabolic intermixing. We considered only the cells which had uniquely-associated ablation marks and excluded 878 cells which were co-sampled (having co-sampling ablation marks, Figure 4H). Still, considering both well-localized metabolites and cells without co-sampling ablation marks only (444 HeLa, 332 NIH3T3) we could predict the cell type with 90.7% accuracy. The lack of the difference in the prediction accuracy (cf. 90.4% for all metabolites and all cells) suggests that co-sampling of cells does not explain the observed metabolic intermixing. In summary, this data indicates that the metabolic intermixing might be a short-distance biological effect when a cell of one type has neighbors of another type in close vicinity and that NIH3T3 cells are stronger affected than HeLa cells.

### SpaceM discovers accumulation of long-chain lipids in steatotic hepatocytes

Next, we used SpaceM to investigate the identity of intracellular metabolomes in differentiated human hepatocytes (dHepaRG). During the non-alcoholic fatty liver disease (NAFLD) hepatocytes are known to accumulate lipid droplets (LDs), which in the context of inflammation and necro-inflammation can lead to development of macrovesicular steatosis (Ringelhan et al. 2018; Wolf et al. 2014). We could recapitulate macrovesicular steatosis using human or murine hepatocytes *in vitro* (Wolf et al. 2014). In particular, the pro-inflammatory cytokine TNFa is known factor promoting steatotic phenotype in hepatocytes (Jung and Choi 2014; Nakagawa et al. 2014). We noticed that the macrovesicular steatosis exhibits a high cell-cell heterogeneity (Figure 5A, Figure S3) and set to characterize the molecular composition of the LDs in steatotic hepatocytes.

**Figure 5.**
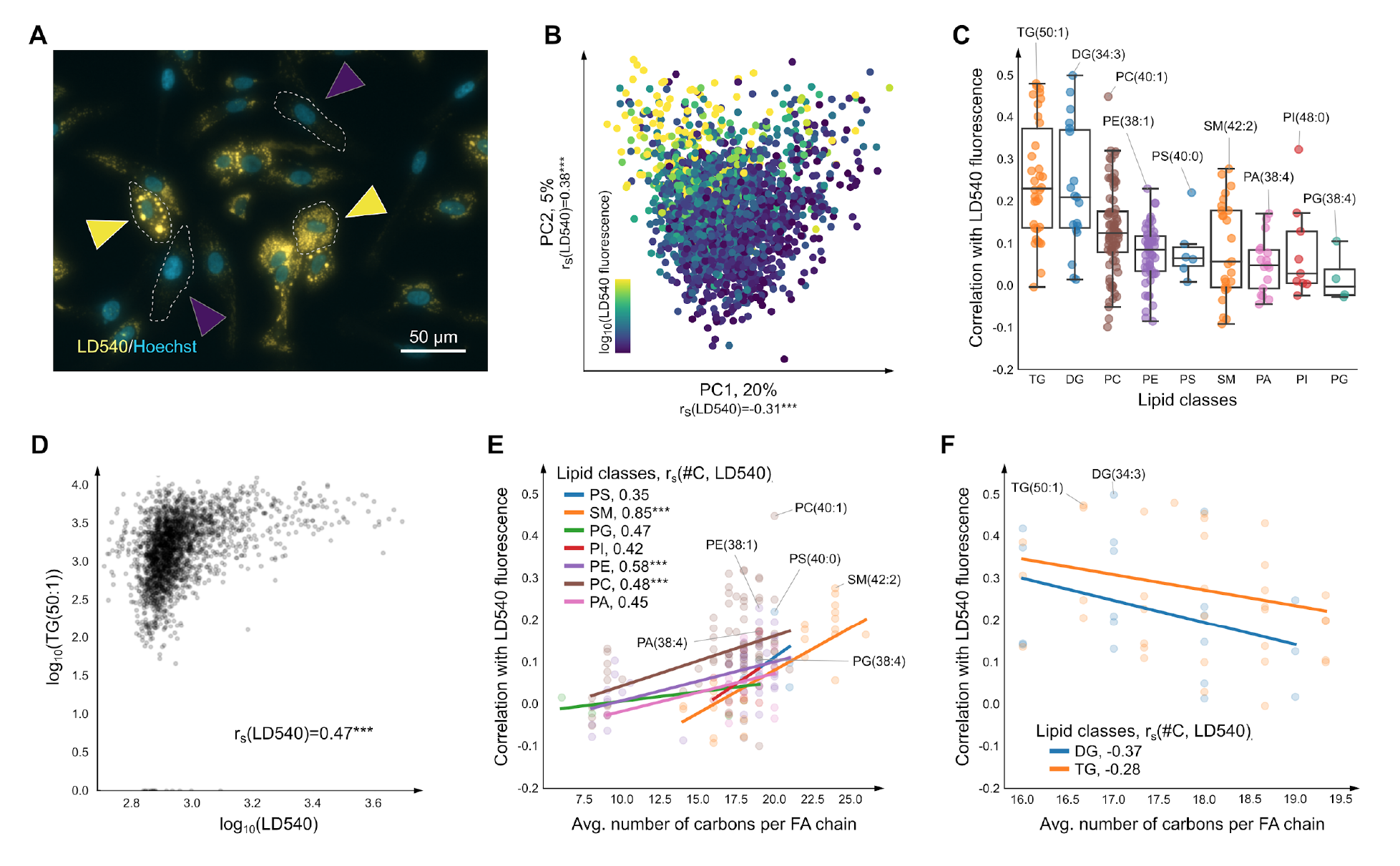
Single-cell analysis of the molecular composition of lipid droplets (LDs) in steatotic hepatocytes stimulated with TNFα (n=2370). **A:** Cell-cell heterogeneity of macrovesicular steatosis (LDs accumulation) in differentiated human hepatocytes stimulated with TNFα in combination with oleic and palmitic fatty acids; the LD540 lipophilic staining highlights intracellular lipid droplets; Hoechst highlights nuclei; the yellow/blue arrows indicate cells with high/low steatosis. **B:** Positive Spearman correlations (rs) the single-cell principal components of the z-scores of the single-cell metabolic profiles between the log10 of LD540 fluorescence prove that the metabolic profiles represent lipid accumulation. **C:** Accumulation of various lipid classes in LDs as measured by the single-cell Spearman correlations between LD540 fluorescence and intensities of 167 detected lipid species; tri- (TG), di-glycerides (DG), and phosphocholines (PC) are the most correlated that is in line with them known to compose the core (TG, DG) and surface (PC) of hepatic LDs. **D:** Single-cell scatterplot of the most-correlated triglyceride TG(50:1), validated using LC-MS/MS (Data S1). **E:** For phosphocholines, sphingomyelins (SMs), and phosphoethanolamines (PEs), the LD540 fluorescence is significantly and positively correlated with the number of carbons (#C) suggesting that steatotic hepatocytes accumulate long-chain versions of these phospholipids; the number of carbons is computed as an average per fatty acid chain to be comparable between lipid classes. **F:** The opposite effect (negative correlation) was observed for di- and triglycerides albeit not significant. *** indicates significance withp-value ≤ 0.001.

We cultured monolayers of differentiated human hepatocytes dHepaRG, stimulated them with TNFa in combination with oleic and palmitic acids, and measured LD accumulation using the lipophilic fluorescent dye LD540 (Spandl et al., 2009). After applying SpaceM, we detected 167 metabolites (at an FDR<10%) in 2370 cells. Principal component analysis (PCA) of the single-cell metabolic profiles revealed a significant correlation between the captured metabolome and the LD540 lipid fluorescence (Figure 5B). Triglycerides (TGs), diglycerides (DGs), and phosphatidylcholines (PCs) were identified as the key constituents of the LDs (Figure 5C). This is in line with neutral lipids, mainly DGs and TGs, known to compose the core of hepatic LDs, whereas polar lipids, primarily PCs, to compose the surface (Gluchowski et al., 2017; Ress and Kaser, 2016). Figure 5D shows a single-cell scatterplot for the most correlated lipid TG(50:1) visualizing statistically significant (with p-value<0.001) correlation of intensities of this lipid with the extent of the macrovesicular steatosis quantified with the LD540 fluorescence. Interestingly, for PCs, sphingomyelins, and phosphoethanolamines, the LD540 fluorescence was found to be positively correlated with the number of carbons suggesting that LDs in steatotic hepatocytes preferentially accumulate long-chain species of these phospholipids (Figure 5E). The opposite effect (negative correlation) was observed for di- and triglycerides albeit not significant (Figure 5F).

### SpaceM discovers intracellular co-regulation of oleic and linoleic acid

We previously found that mice fed a diet enriched with oleic and palmitic fatty acids developed key features of human metabolic syndrome, nonalcoholic steatohepatitis (NASH), and NASH-derived hepatocellular carcinoma (Wolf et al., 2014). To investigate on the single-cell level how the metabolome and lipidome of human hepatocytes is affected by different pro-inflammatory factors, we analyzed hepatocytes cultured under the following conditions: (i) CTRL, untreated cells, (ii) FA, cells stimulated with oleic and palmitic fatty acids (opFAs), (iii) LPS, cells stimulated with a pathogen-associated molecular pattern lipopolysaccharide and opFAs and (iv) TNFa, cells stimulated with TNFα and opFAs. SpaceM obtained metabolic profiles of 136 metabolites for 22258 cells in total with a batch correction applied to three randomized technical replicates per condition (Table S1, Figure S4). LD540 staining of differentiated, stimulated human hepatocytes revealed lipid accumulation and macrovesicular steatosis (Figs. 5A, S6, S8). PCA of single-cell metabolic profiles indicates the captured differences in the metabolomes of untreated and stimulated cells (Figure 6A). The metabolic shifts between different conditions reflected the expected levels of response to the stimuli as (i) opFAs used for stimulation in the FA condition were also supplemented in the LPS and TNF condition, (ii) TNFa specifically induced the strongest TNF-receptor signaling whereas (iii) LPS only secondarily induces TNF secretion and TNFR signaling at lower levels (Beutler, 2004). We investigated the contributions of individual metabolites to the principal components and found that, as expected, cells cultured with oleic acid accumulated intracellular oleic acid as compared to the untreated cells (Figure 6B). Not all molecules showed elevated accumulation with the increase of the stimuli; see e.g. PIP(38:5) in Figure 4C which exhibits similar intensities between CTRL and FA with a clear increase in the TNF and even more in the LPS condition. Interestingly, opposite to oleic acid, linoleic acid (the second of the opFAs stimuli in the FA, LPS, TNF conditions) exhibits decreased levels in the stimulated cells (Figure 6D). Importantly, there is a clear correlation between the levels of the oleic and linoleic acids across all cells (Figure 6E). A similar effect of the inverse relation between oleic and linoleic acids levels was reported in the livers of mice fed a high-fat diet (da Silva-Santi et al., 2016). However, the bulk analysis could not discern whether the effect occurs in different cell subpopulations or is concerted within the same cells. Our single-cell analysis shows that the effect happens indeed within the same cells thus suggesting intracellular co-regulation of oleic and linoleic acid levels.

**Figure 6.**
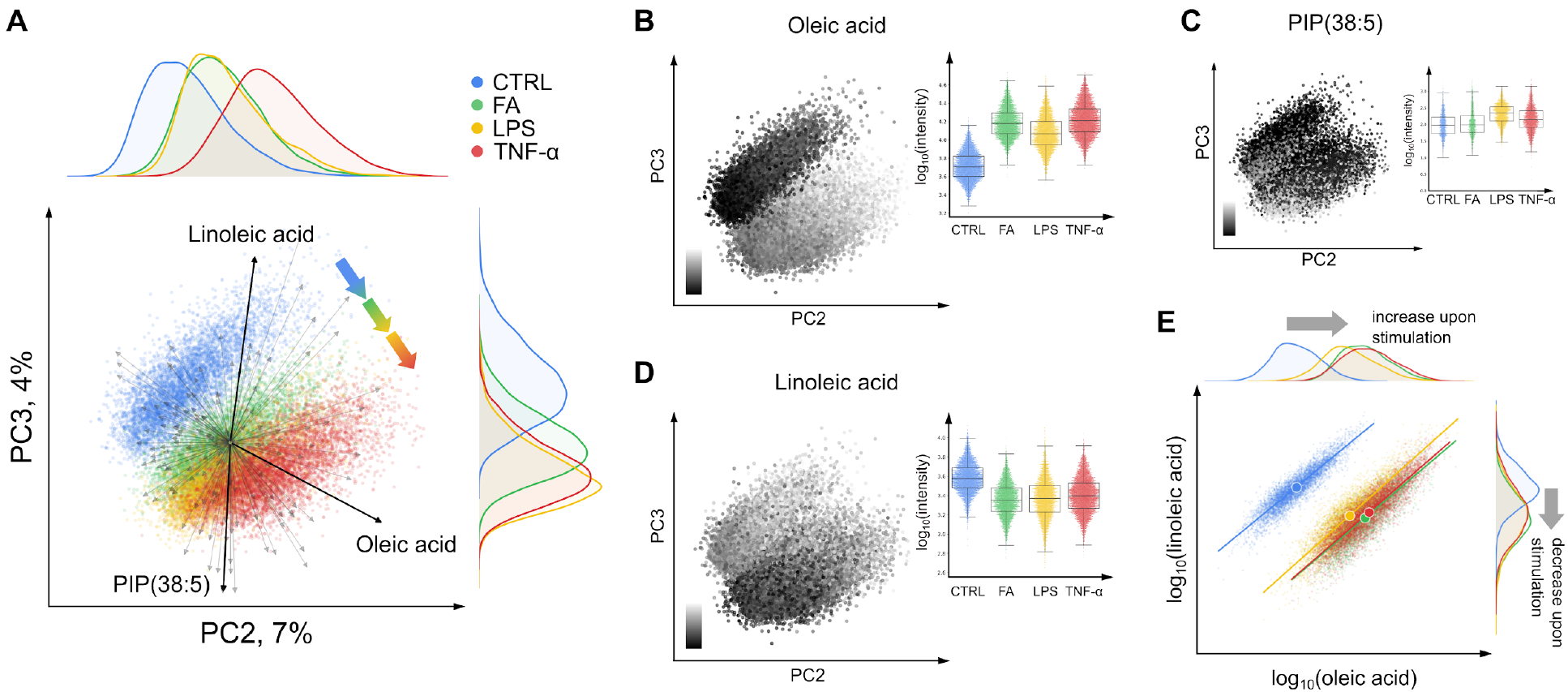
Single-cell statistical analysis of steatotic differentiated human hepatocytes dHepaRG stimulated with the fatty acids, LPS, and TNFα (n=22258). **A:** PCA of z-scores of the single-cell profiles of 136 metabolites; biplot vectors visualize contributions of individual metabolites; gradient-colored arrows illustrate the metabolomes transitions from the untreated cells (CTRL, n=5654) to the cells cultured with oleic and palmitic fatty acids (FA, n=4972), with LPS and the fatty acids (LPS, n=5062), or with TNFα and the fatty acids (TNFα, n=6570). **B-D:** Single-cell metabolite intensities mapped onto the PCA plot and the Tukey box plots per condition (25%-75% percentiles, whiskers at 1.5x distance of the interquartile range); only cells with non-zero metabolite intensity are shown; see Figure 14 for AMP and glutathione. **E:** Single-cell scatterplot for intensities of linoleic vs. oleic acids showing an inverse relationship in intracellular levels of these fatty acids upon stimulation and their tight and condition-independent correlation; the centers of masses and fitted lines are plotted. Oleic acid, linoleic acid, and PIP(38:5) were validated using LC-MS/MS (Data S1).

### SpaceM discovers association of PIP(38:5) with high cell-cell contact

Finally, we investigated the spatio-molecular organization of human hepatocytes (Figure 7A-D). LD540-fluorescence microscopy revealed lipid accumulation within groups of cells that display high cell-cell contact (Figure S5, S6). Among all the detected metabolites, phosphatidylinositol phosphate PIP(38:4) (although not validated with LC-MS/MS) and PIP(38:5) were most highly associated with cell-cell contact (Figure 7B, Figure S7). PIPs are precursors of PIP3, a signaling phospholipid in the plasma membrane, which might explain their elevation within adjacent cells in a locally-concerted manner. Not all detected metabolites displayed such an association: AMP showed no correlation (Figure 7C) whereas oleic acid showed a negative correlation (Figure 7D). Thus, integrating fluorescent, morphological, spatial, and molecular information can be a powerful approach to explore multi-cellular phenomena (Bray et al., 2016).

**Figure 7.**
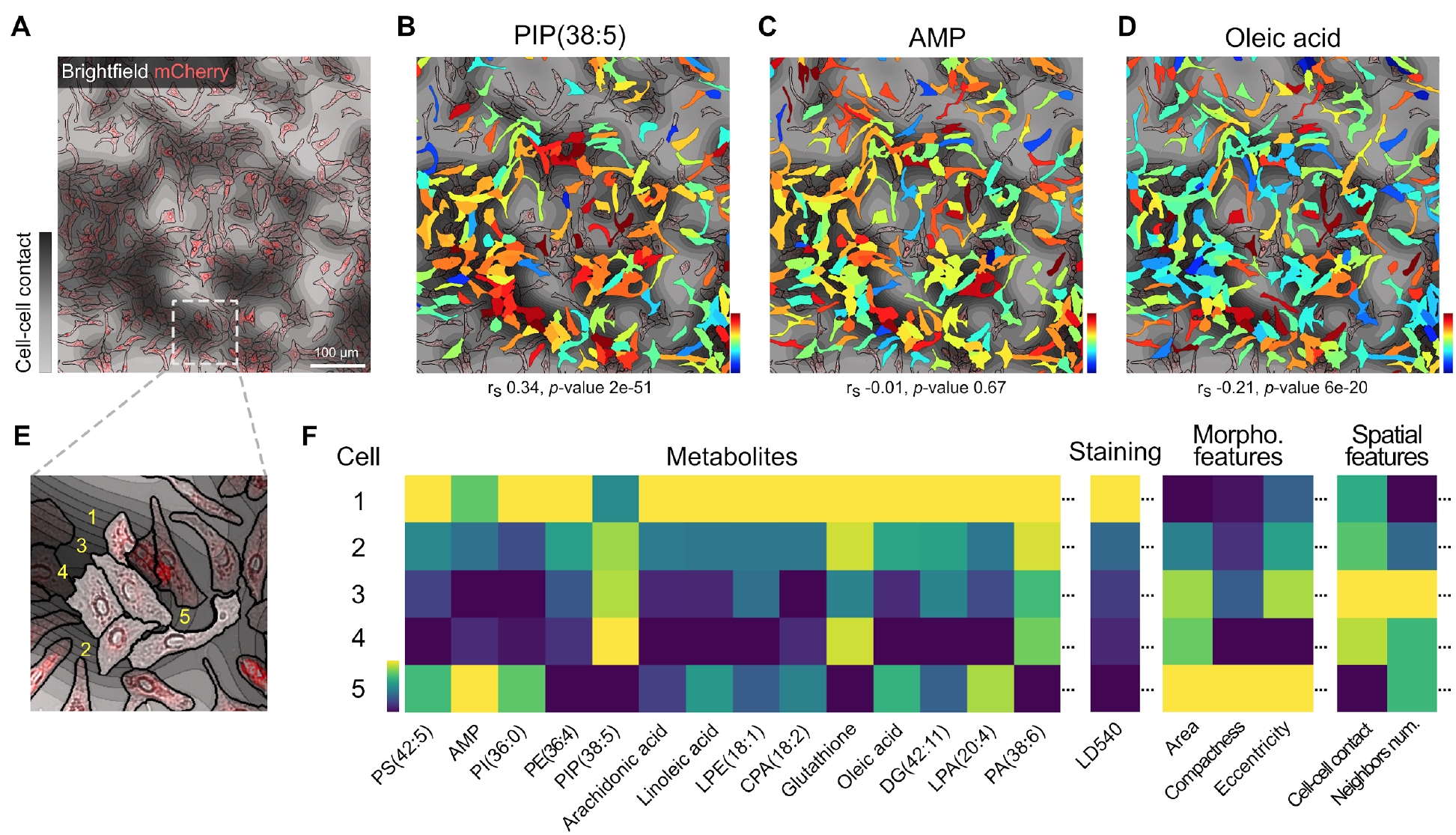
Single-cell statistical analysis of steatotic differentiated human hepatocytes stimulated with pro-inflammatory factors (n=22258). **A:** TNFα hepatocytes with areas of local crowding; the red LD540 fluorescence intensity indicates the accumulation of lipid droplets from low steatosis to macrovesicular steatosis. **B-D:** Single-cell molecular images for PIP(38:5), AMP, and oleic acid. Next to PIP(38:4), PIP(38:5) is the second most correlated molecule with cell-cell contact, indicating its potential concerted elevation within adjacent cells. Oleic acid, linoleic acid, AMP, glutathione, and PIP(38:5) were validated using LC-MS/MS (Data S1). **E-F:** Illustration of a part of spatio-molecular matrix for five selected cells that represents single-cell phenotypic information (LD540 fluorescence quantifying the extent of the macrovesicular steatosis), morpho-spatial features (including cell-cell contact), and metabolite intensities.

## Discussion

Here, we presented SpaceM, a novel method for the spatial single-cell analysis of intracellular metabolomes of cultured cells. SpaceM not only detects metabolites in individual cells using MALDI but, critically, integrates MALDI analysis with microscopy and is supplied with data analysis strategies for metabolite annotation, intensities normalization, selection intracellular metabolites and filtering out poor quality cells. The method demonstrated to be robust and reproducible, allowing us to analyze several conditions and replicates, obtaining metabolic profiles of around 100 metabolites from over 30.000 cells.

We benchmarked the method by analysing the co-cultured human HeLa cells and mouse NIH3T3 fibroblasts. The single-cell metabolic profiles detected by SpaceM were rich enough to predict the cell type with 90.4% accuracy with the single-cell resolution. Surprisingly, we detected metabolic intermixing of different cell types upon co-culturing, a yet unreported biological phenomenon. Our investigations indicated that the observed intermixing is a short-distance effect, namely, its extent depends on the presence of cells of other type in close vicinity. We carefully considered all other confounding factors and outruled sampling inaccuracies or metabolite delocalization. All our results indicate that intracellular metabolomes of cells of one type are indeed influenced by the neighboring cells of another type, with NIH3T3 cells affected stronger than HeLa cells.

For molecular detection, SpaceM exploits MALDI-imaging thus inheriting the advantages of this technique, in particular the capacity for untargeted metabolomics. We demonstrated it by detecting profiles encompassing 88 (for HeLa and NIH3T3) and 136 (for dHepaRG) metabolite annotations on the level 2 according to the Metabolomics Standards Initiative (Sumner et al., 2007). The use of METASPACE for metabolite annotation was instrumental in the interpretation, quality control, and fast access to the metabolite images of the collected MALDI-imaging data, as METASPACE efficiently reduces millions of mass-to-charge values to tens of metabolite annotations in a False Discovery Rate-controlled manner. SpaceM is not limited to lipids, fatty acids and such small molecules as AMP or glutathione (Figure S8, Data S1) but can be extended to other molecular classes by using another MALDI-imaging protocol. In the HeLa-NIH3T3 co-culture benchmarking experiment, even when using just a half of the 88 detected metabolites, we could predict the cell type with 87.6% accuracy that affirms the richness of the metabolic profiles detected by SpaceM.

SpaceM was enabled by MALDI-imaging achieving the single-cell spatial resolution. However, it is almost impossible to sample highly-confluent cell monolayers without co-ablating several cells at once. Moreover, the sample preparation for MALDI-imaging, particularly matrix application, can negatively affect the spatial resolution. Thus, we set to prove the single-cell nature of the method, especially because it was essential to exclude possible technical reasons for the observed metabolic intermixing between HeLa and NIH3T3. For this, we developed a strategy to consider for each metabolite its delocalization outside the cell perimeter. This led to revelation that, first, the extent of delocalization is metabolite-specific, and second, even for the minority of metabolites characterized as delocalized, the median delocalization is comparable to the average size of a single cell. Moreover, in our benchmarking experiment, discarding delocalized metabolites did not affect the accuracy of the cell type prediction much. Altogether, these novel delocalization analyses confirm the single-cell nature of the SpaceM method. We hypothesize that the delocalization happens during spraying the MALDI matrix solution but do not have yet a definite answer what makes some metabolites delocalized while the majority of metabolites were detected well-localized within the cell perimeter. For example, lysophosphoethanolamine LPE(18:1) was found to be well-localized whereas the structurally similar but saturated LPE(18:0) was found to be delocalized.

Compared to other reported single-cell metabolomics methods, e.g. microwells (Ibáñez et al., 2013) or microsampling (Guillaume-Gentil et al., 2017) approaches, SpaceM analyzes cells *in situ* in their native spatial context, ensures minimal unwanted perturbation, and preserves information about the microenvironment and spatial organization of cells. In contrast to micro-and nano-sampling methods, SpaceM is also a high-throughput method able to analyze over 10000 cells and at the same time, as we illustrated, detecting rich metabolic profiles. Compared to a microscopy-guided laser ablation approach (Do et al., 2017), SpaceM uses unbiased sampling that facilitates discovery of cell populations which cannot be discriminated by microscopy, helps distinguish intracellular from extracellular signals, and also capitalizes on a softer MALDI ionization better suited for biomolecules. Compared to ultra-high spatial resolution approaches (Passarelli et al., 2017), SpaceM makes possible a high throughput analysis of large populations of cells to investigate their heterogeneity and to discover rare molecular phenotypes. The combination of these strengths makes SpaceM not only a single-cell but also a spatial method. We demonstrated the spatial capacity of SpaceM by discovering short-distance effect of metabolic intermixing between HeLa and NIH3T3 cells and by associating PIPs with high cell-cell contact.

We expect SpaceM to be broadly applicable to any adherent cells cultured in a monolayer, avoiding growing on top of each other that can lead to increased co-sampling. In our experience, cell culturing for SpaceM is relatively straightforward and can be evaluated following conventional cell biology practices by paying attention to the cell count, viability and confluence. SpaceM allows for the determination of cells that are different in their response to changes in the microenvironment, which enables the identification of novel molecular mechanism involved in critical biological processes.

SpaceM contributes to the growing field of single-cell-omics methods by providing the missing capacities for spatio-molecular *in situ* analysis. Future experiments will aim to translate SpaceM to the level of tissue sections. Our method will be particularly useful to investigate health and disease phenomena associated with metabolic reprogramming, spatial organization and/or cellular heterogeneity such as differentiation, infection, drug metabolism, immunity, and cancer.

## Acknowledgements

We thank Nassos Typas for advising on biology and providing access to the microscope, Bashir El Debs and Joel Selkrig for training on the microscopy and cell culturing (all EMBL), Andrew Palmer (EMBL) for training and support on MALDI-imaging, Megan Stanifer and Steeve Boulant (DKFZ) for their support and training on the cell culturing, METASPACE software development team (EMBL) for creating and supporting the METASPACE software, Carina Beatrice Vibe (EMBL) for her support and feedback on the manuscript, Angela Andersen (Life Science Editors) for scientific editing, Samantha Seah and Christoph Merten (EMBL) for providing the NIH3T3-GFP cell line, Fabian Merkel and Christian Häring (EMBL) for providing the HeLa Kyoto H2B-mCherry cell line. We further thank other members of the Thesis Advisory Committee of L.R.: Anne-Claude Gavin (EMBL) and Britta Brügger (Heidelberg University). This work was supported by the European Union’s Horizon 2020 program under the grant agreements 634402 (T.A.) and 667273 (M.H.), the DKFZ-MOST cooperation program (M.H., M.S.), Darwin Trust of Edinburgh (S.T.), SFB Transregio grants 179, 209, and 1335 (all M.H.), and the ERC Consolidator grants HepatoMetaboPath (M.H.) and METACELL (T.A.).

## Author Contributions

L.R. and T.A. conceived the research. L.R. developed the method. T.A. supervised the study. S.T. performed the co-culture experiment. P.P. performed LC-MS/MS validation. M.S. and M.H. contributed with the hepatocytes model, conceived the treatment design. M.S. cultured and prepared hepatocytes. L.R., M.S., M.H. and T.A. interpreted data. L.R. and T.A. wrote the paper.M. S. and M.H. contributed to the paper writing.

## Methods

### Co-culturing of HeLa and NIH3T3 cells

HeLa Kyoto H2B-mCherry and NIH3T3-GFP cells were cultured at 37 °C with 5% CO2, and were maintained in high glucose DMEM (1X Pen/Strep) (Gibco/ThermoFisher Scientific, Bremen, Germany) supplemented with 10% FBS, 100 U/ml penicillin, 100 µg/ml streptomycin (Gibco) and 1 mM sodium pyruvate (Gibco). Cells were trypsinized with 0.25% trypsin-EDTA (Gibco) and split 1:10 twice a week. Two technical replicates for the co-cultures and one replicate for monoculture were used. Trypsinized cells were counted and cells were seeded on 4- well-glass labtek chamber slides (Lab-Tek II, CC2) (ThermoFisher Scientific). In the co-culture experiment, equal number of cells of each cell type was added into each well (4×10^5^ cells/well). After 48h of incubation cells were washed with PBS. After washing, the cells were fixed for 15 min with 4% paraformaldehyde (Sigma Aldrich, Darmstadt, Germany) at room temperature. Then the cells were stained with DAPI (1µg/ml) (ThermoFisher Scientific) in PBS for 20 min at room temperature.

### Hepatocytes culturing and stimulation

HepaRG cell culture and differentiation was performed as described earlier (Gripon et al., 2002). Differentiated HepaRG (dHepaRG) cells were cultured on 4-well-glass chamber slides (Lab-Tek II, CC2, ThermoFisher Scientific, Bremen, Germany) (5.5×10^4^ cells/well). The cells were stimulated with the fatty acids (opFAs): oleic acid (66 µM) and palmitic acid (33 µM), opFAs and tumor necrosis-alpha (TNFα) with the final concentration of 50 ng/ml (Recombinant Human TNF-alpha, and Systems), or opFAs and lipopolysaccharide (LPS) (100 ng/ml) (LPS from *E.coli*) (Sigma Aldrich) in Williams E Medium (William’s Medium E, with stab. glutamine, without Phenol Red, with 2,24 g/l NaHCO3) (PAN Biotech) for 24 h. Cells grown in Williams E Medium without supplement for 24 h were considered as a negative control. For each of those four conditions, cells were seeded in three different wells which were considered as technical replicates (Table S1). After washing, cells were fixed for 15 min with 4% paraformaldehyde (Sigma Aldrich) at room temperature. Then the cells were washed and stained with Hoechst (1µg/ml) (Hoechst 33342) (ThermoFisher Scientific) and LD540 (0.1 µg/ml) (Spandl et al., 2009) in PBS for 30 min at room temperature. After washing, cells were stored in dH2O at 4 °C for one night maximum.

### Preparing cells for imaging

The plastic walls of the labtek were removed and the cells were dried in a Lab Companion^TM^ Cabinet Vacuum Desiccator for 16h at room temperature and −0.08 MPa. After complete desiccation of the cells, pen marks are manually drawn on the glass slide using a black alcohol pen model 140s black (Edding, Ahrensburg, Germany) to keep track of the glass slide orientation and for image registration. The marks were drawn on the same side as the cells. Cells are kept at 4 °C until analysis. For the following experiments, the samples were analyzed by the microscopy and MALDI-imaging mass spectrometry following a randomized experimental design (Table S1).

### Pre-MALDI bright-field and fluorescence microscopy of cells

Fixed cells were sequentially observed with the camera Nikon DS-Qi2 (Nikon Instruments) with the Plan Fluor 10x (NA 0.30) objective (Nikon Instruments) mounted on the Nikon Ti-E inverted microscope (Nikon Instruments) in bright-field and fluorescence (620 nm and 460 nm). The pixel size was 0.73 µm. The microscope was controlled using the Nikon NIS Elements software. The tiled acquisition of each cell culture area was performed using the JOB functionality of the NIS software. Stitching of tiled frames was performed using the FIJI stitching plugin (Preibisch et al., 2009).

### MALDI imaging mass spectrometry

Relative humidity and temperature levels in the mass spectrometry room were monitored and controlled during the whole experiment and were within 44%-63% and 21.1-23.7 °C (Table S1). For the analysis of the lipid droplets (Figure 3), the 2,5-dihydroxybenzoic acid (DHB) matrix (Sigma Aldrich) 15mg/ml dissolved in 70% acetonitrile was applied onto the dried cells on the labtek slides by using a TM-Sprayer robotic sprayer (HTX Technologies, Carrboro, NC, USA). Spraying parameters were as following: temperature=100 °C, number of passes=8, flow rate=0.07 ml/min, velocity=1350 mm/min, track spacing=2 mm/min, pattern=CC, pressure=10 psi, gas flow rate=5 l/min, drying time=15 sec, nozzle height=41 mm. The estimated matrix density was of 0.00311 mg/mm^2^. For investigating the molecular trends within all four considered conditions (Figure 4), the matrix 1,5-diaminonaphthalene (DAN) (Sigma Aldrich) 10mg/ml dissolved in 70% acetonitrile was applied onto the dried cells on the labtek slides by using the same TM-Sprayer robotic sprayer. Spraying parameters were as following: temperature=90 C°, number of passes=8, flow rate=0.07 ml/min, velocity=1350 mm/min, track spacing=3 mm/min, pattern=CC, pressure=10 psi, gas flow rate=2 l/min, drying time=15 sec, nozzle height=41 mm. The estimated matrix density was of 0.001383 mg/mm^2^. For MALDI imaging mass spectrometry, the glass slides with the dried cells on them were mounted onto a custom slide adaptor and loaded into the AP-SMALDI source (Transmit, Giessen, Germany). The MALDI laser focus was optimized manually using the source cameras with the focused beam diameter estimated to be between 15.0 and 43.0 µm (mean equal to 29.9 µm, standard deviation equal to 8 µm). The x-y step size (distance between the centers of ablation marks) was set to 50 µM. For each pixel, the spectrum was accumulated from 30 laser shots at 60 Hz. Negative mode MS analysis was performed in the full scan mode in the mass range of 200-1100 m/z (resolving power R=140000 at m/z 200) using an QExactive Plus mass spectrometer (ThermoFisher Scientific). MS parameters in the Tune software (version 2.5 Build 2042, ThermoFisher Scientific) were set to the spray voltage of 4.10 kV, S-Lens 80 eV, capillary temperature 250 C. The data was converted from the RAW format into the imzML format containing only centroided data using the ImageQuest software, v.1.1.0 (ThermoFisher Scientific). Metabolite annotation was performed using the METASPACE cloud software (http://metaspace2020.eu) implementing the bioinformatics methods for False Discovery Rate-controlled annotation published by us earlier (Palmer et al., 2017) with the m/z tolerance of 3 ppm and FDR of 10%, 20%, and 50% against the HMDB metabolite database v2.5 (Wishart et al., 2009).

### Post-MALDI microscopy to detect MALDI ablation marks

The cells were imaged in bright-field microscopy after MALDI-imaging using the same microscopy setup and parameters as described earlier in the pre-MALDI microscopy section to define the positions of the ablation marks with respect to the fiducial marks.

### Association of laser ablation marks with single cells

This is the key part of the method as it solves the challenge that single cells are not visible in the post-MALDI microscopy images due to the opaque layer of MALDI matrix covering cells. Here, ablation marks left by the MALDI laser were associated with single cells in three steps: a) cells segmentation in the pre-MALDI microscopy images, b) detection of laser ablation marks in post-MALDI microscopy images, c) matching between ablation marks and MALDI mass spectra and d) co-registration of pre- and post-MALDI microscopy images to overlay the ablation marks with the segmented single cells.

In step a), cells were segmented using a custom pipeline in the CellProfiler software (Carpenter et al., 2006) where the DAPI staining channel was used to generate seeds for a region growing algorithm detecting cells boundaries in the LD540-staining channel. In step b), we first denoised the bright-field microscopy images by applying a low-pass filter in the 2D Fourier frequency domain, in particular to exploit both the regular distances between ablation marks as well as the repeated shape of the ablation mark itself. Then, we applied a contrast-enhancing filter (using the *clip* function from the Python module *numpy*) and Otsu’s thresholding method (Otsu, 1979) to binarize the image (using the *imbinarize* function in Matlab). Then, we applied morphological image analysis operations of closing and then opening to fill in the holes in the image and to remove individual noisy pixels (using the *imclose* and *imopen* functions in Matlab). This provided estimations of the centers of mass of each ablation mark (Figure S9). In step c), we fit a theoretical rectangular grid to the ablation marks. The numbers of X- and Y- grid steps were defined as set up during the MALDI acquisition. The center of the acquisition region was considered as the center of the grid. The orientation of the grid with respect to the post-MALDI microscopy image was optimized by finding an angle which resulted in the best overlap between the grid lines and the detected ablation marks. The X- and Y-spacing of the grid were optimized by minimizing the distance between the grid nodes and the center of mass of the nearest neighbor ablation mark. Then, only ablation marks which were the nearest neighbors to the grid nodes were taken and re-indexed (Figure S10). This provided X- and Y-coordinates for each ablation mark associated with a collected MALDI spectrum. In order to improve estimations of the ablations marks areas used later for normalization, their segmentation was further improved by applying a custom region-growing algorithm implemented in Python. In step d), co-registration of pre- and post-MALDI microscopy images was done based on the pen marks drawn on the edge of the wells used as fiducials. We first segmented the pen marks in both pre- and post-MALDI bright-field microscopy images using Otsu’s intensity thresholding method. Then, we used the basin-hopping optimization algorithm (Python implementation from the *scipy* package v0.18.1) to find the best linear transformation matching the coordinates of the edges of the pen marks between the pre- and post-MALDI images (Figure S10). The optimal linear transformation was applied to the post-MALDI microscopy images to map the ablation marks to the pre-MALDI microscopy images. The initial assessment of the co-registration quality and overlaying of the metabolite intensities was performed in the ‘ili web app at http://ili.embl.de (Protsyuk et al., 2018).

### Single-cell intensity normalization

A normalized intensity of each metabolite in a single cell was constructed as follows. For each cell, we considered all ablation marks overlapping with the cell area and selected the associated ablation marks which overlap with the cell by over than 30% of their ablation area. The metabolite intensities coming from an ablation mark were normalized by dividing them by the ratio of the sampling area (defined as the number of pixels of the intersection of the ablation mark and any cell region) to the area of the ablation mark. Finally, for each cell its normalized metabolite intensities were calculated as the weighted average normalized intensities of the associated ablation marks where the weights are defined as the ratio of the shared pixels (Figure 2). In order to account for the variations in permeabilization efficiency between the biological replicates, single-cell LD540 fluorescence intensities were normalized by dividing them by the median DAPI intensity (median over a well).

### Selecting intracellular metabolites

We selected metabolite annotations corresponding to intracellular metabolites as follows. First, for each ablation mark we assigned to it the inside-cells label having values either of zero or one based on whether the mark has any overlap with any cell. Then, for each metabolite ion image, its intensities were binarized to zero-one values by selecting a threshold leading to the highest Pearson correlation with the inside-cells labels. The threshold value was found using the basin-hopping optimization algorithm. In order to consider only intracellular metabolites for further analysis, we selected those metabolite annotations whose binarized ion images were correlated with the inside-cells labels with the Pearson correlation higher than 0.25. For the stimulated dHepaRG experiment where three replicates for each of four conditions were obtained, we considered the metabolite annotations which were shared by at least three samples (out of 12 overall) that led to 136 annotations. For each metabolite annotation, we pulled the ion images with the m/z tolerance of 3ppm from the imzML files.

### Cell filtering and batch correction

We filtered out 5% of cells with the lowest metabolite yield, namely the cells which had most zero-valued metabolites annotations, following the approach well-accepted in single-cell transcriptomics (Grün and van Oudenaarden, 2015) (Figure S11). In the stimulated HepaRG experiment, this filtered out 1240 cells out of 23498 overall. In the stimulated HepaRG experiment, to compensate for the batch effect between the biological replicates within each condition, we applied the *combat* batch correction algorithm (Fortin et al., 2017) originally developed for single-cell transcriptomics data using its open-source Python implementation neuroCombat available at https://github.com/ncullen93/neuroCombat (Figure S4).

### Cell type classification for the co-culture experiment

The assignment of the cell type based on the constitutive fluorescence of the cells (mCherry for HeLa, GFP for NIH3T3) was done by finding a separating linear boundary between the two populations (Figure 2A). The resulting cell types provided the ground truth labels for the Linear Discriminant Analysis (LDA) performed using the Python *scikit-learn* LDA implementation (version 0.19.1).

### LC-MS/MS validation of METASPACE annotations

#### Sample preparation

Lipids and fatty acids were extracted using the Folch method (Folch et al., 1957) with chloroform:methanol (2.5:1). For lipidomics analysis, the dried samples were reconstituted in isopropanol:methanol (1:1) and injected 10uL into the LC-MS system. For metabolomics analysis, metabolites were extracted in 80% methanol and directly injected 20uL into the LC-MS system.

#### LC-MS/MS methods for lipidomics

All LC-MS/MS analyses were performed on a Vanquish Ultra-High Performance Liquid Chromatography (UHPLC) system coupled to a Q-Exactive Plus High Resolution Mass Spectrometry (HRMS) (ThermoFisher Scientific) with an electrospray ionization (ESI) source operated in either positive or negative mode. The separation of lipids and fatty acids was carried out using an Agilent Poroshell EC-C18 column (3 x 50 mm; 2.7 µM) maintained at 40 °C at the flow rate of 0.26 ml/min. The mobile phase consisted of solvent A (acetonitrile-water (4:6)) and solvent B (isopropyl alcohol-acetonitrile (9:1)), which were buffered with either 10 mM ammonium acetate (for negative mode) or 10mM ammonium formate acidified with 0.1% formic acid (for positive mode). The UHPLC gradient was set at 20%, 20%, 45%, 52%, 66%, 70%, 75%, 97%, 97%, 20%, 20% of solvent B at the time points 0, 1.5, 4, 5, 7, 8, 10, 12, 15, 16, 19 min, respectively. Fatty acids and lipids were detected with the HRMS full scan at the mass resolving power R=35000 in the mass range of 100-1500 m/z. The data-dependent tandem (MS/MS) mass scans for five most intense ions (TOP5) were obtained along with full scans using higher energy collisional dissociation (HCD) with normalized collision energies of 20, 30 and 40 units at the mass resolving power R=17500. The MS parameters in the Tune software (ThermoFisher Scientific) were set as: spray voltage of 4 kV, sheath gas 30 and auxiliary gas 5 units, S-Lens 65 eV, capillary temperature 320 ^o^C and vaporization temperature of auxiliary gas was 300 ^o^C.

#### LC-MS/MS methods for metabolomics

LC-MS/MS metabolomics analysis was carried out using an Xbridge BEH Amide column (100X 2.1 mm; 2.5 µM) maintained at 40 °C at the flow rate of 0.3 ml/min. The mobile phase consisted of solvent A (7.5 mM ammonium acetate with 0.05% NH4OH) and solvent B (acetonitrile). The UHPLC gradient was set at 85%, 85%, 10%, 10%, 85%, 85% of solvent B at the time points 0, 2, 12, 14, 14.1, 6 min, respectively. Metabolites were detected with HRMS full scan at the mass resolving power R=70000 in mass range of 60-900 *m/z*. The data-dependent MS/MS mass scans were obtained along with full scans using HCD of normalized collision energies of 10, 20 and 30 units which were at the mass resolving power R=17500. The MS parameters in the Tune software (ThermoFisher Scientific) were set as: spray voltage of 4 kV (for negative mode 3.5 kV), sheath gas 30 and auxiliary gas 5 units, S-Lens 65 eV, capillary temperature 320 ^o^C and vaporization temperature of auxiliary gas was 300 ^o^C. Data was acquired in the full scan mode and MS/MS mass spectra for TOP5 precursor ions.

#### LC-MS/MS validation of METASPACE annotations

LC-MS/MS validation of lipid and metabolite METASPACE annotations was performed either by comparing retention times, exact m/z (MS) and fragmentation pattern (MS/MS) spectra with authentic standards or by matching MS/MS spectra with the EMBL Metabolomics Core Facility (MCF) spectral library (available at http://curatr.mcf.embl.de/) and public spectral libraries (LipidBlast, LIPID MAPS and mzCloud). The details of annotation validation are summarized in Supplementary Data S1. The structural annotation procedure for head groups (HD) and fatty acid side chains (SD) is described in details in (Palmer et al., 2017).

### Data visualization

All plots were generated in Python, version 3.6.2, by using the packages *matplotlib* 2.1 and *seaborn* 0.8.1. The Python package *scikit-learn* 0.19.1 was used for the Principal Component Analysis.

### Data availability

All metabolite and lipid annotations and images are publicly available at the METASPACE online knowledgebase (URL for the co-cultured and mono-cultured HeLa and NIH3T3 cells from Figure 2, URL for the dHepaRG cells from Figure 3, URL for the dHepaRG cells from Figure 4).

## Supplementary Figures

**Supplementary Figure S1.**
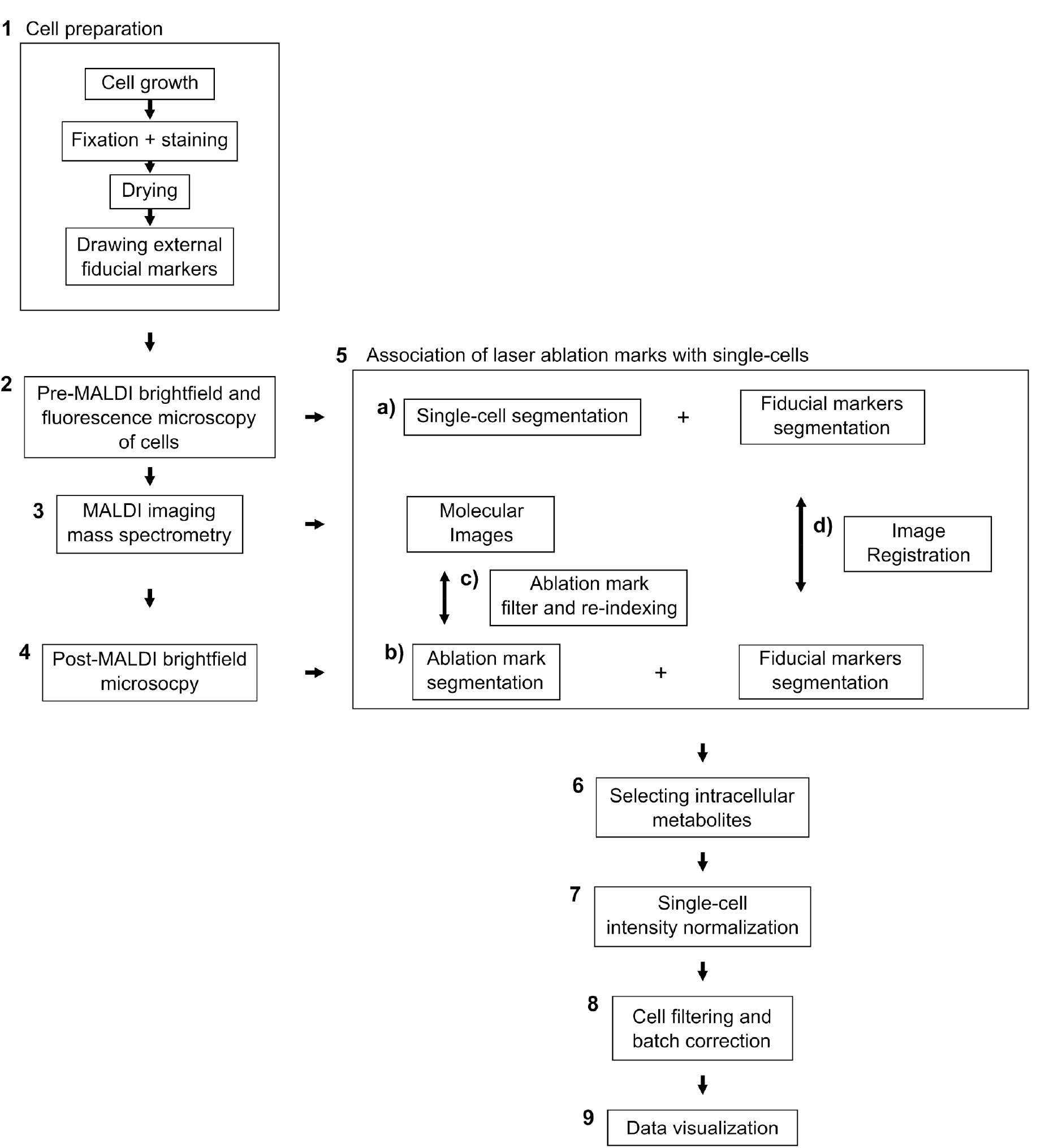
Detailed workflow of the SpaceM method, see Figure 1 for a visualization of the workflow supplemented with visual elements.

**Supplementary Figure S2.**
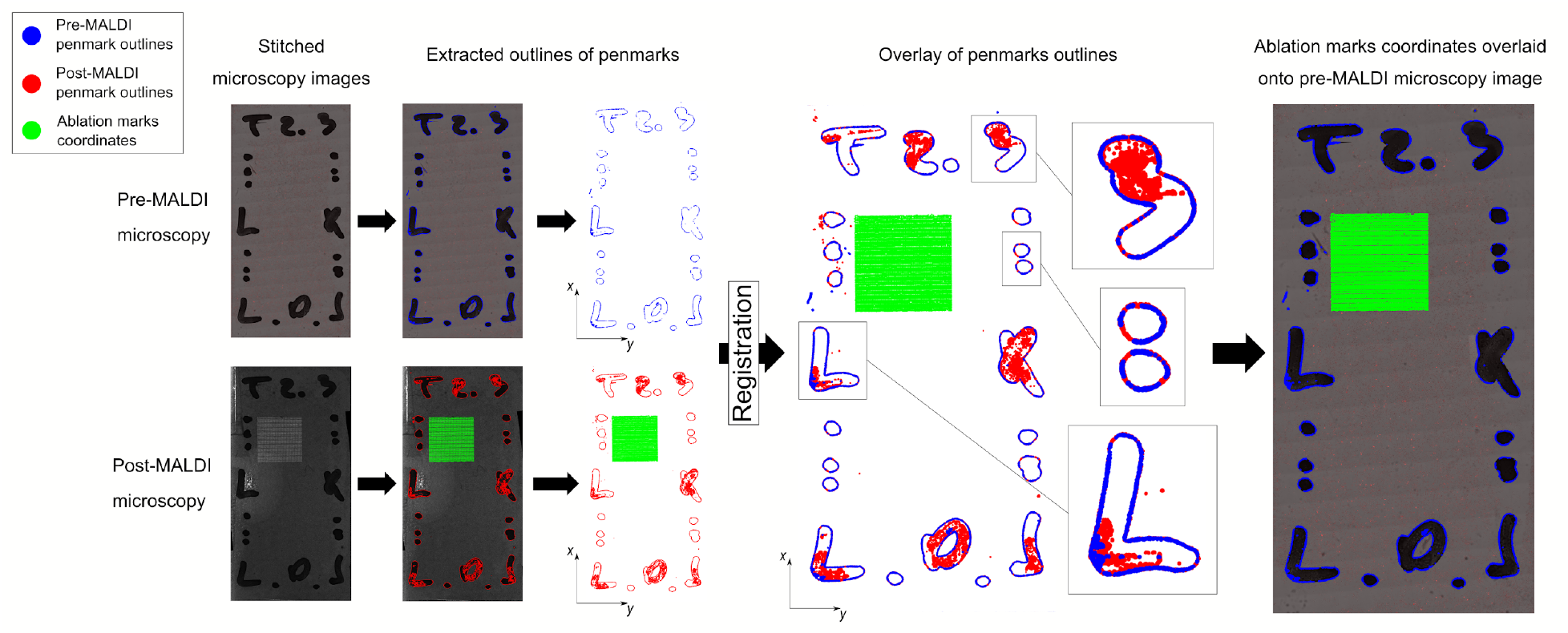
Registration workflow of pre- and post-MALDI microscopy images. The area where MALDI-imaging was applied is shown in green. The features of the pen marks used as fiducials are shown in blue (for pre-MALDI microscopy images) or red (for post-MALDI microscopy images).

**Supplementary Figure S3.**
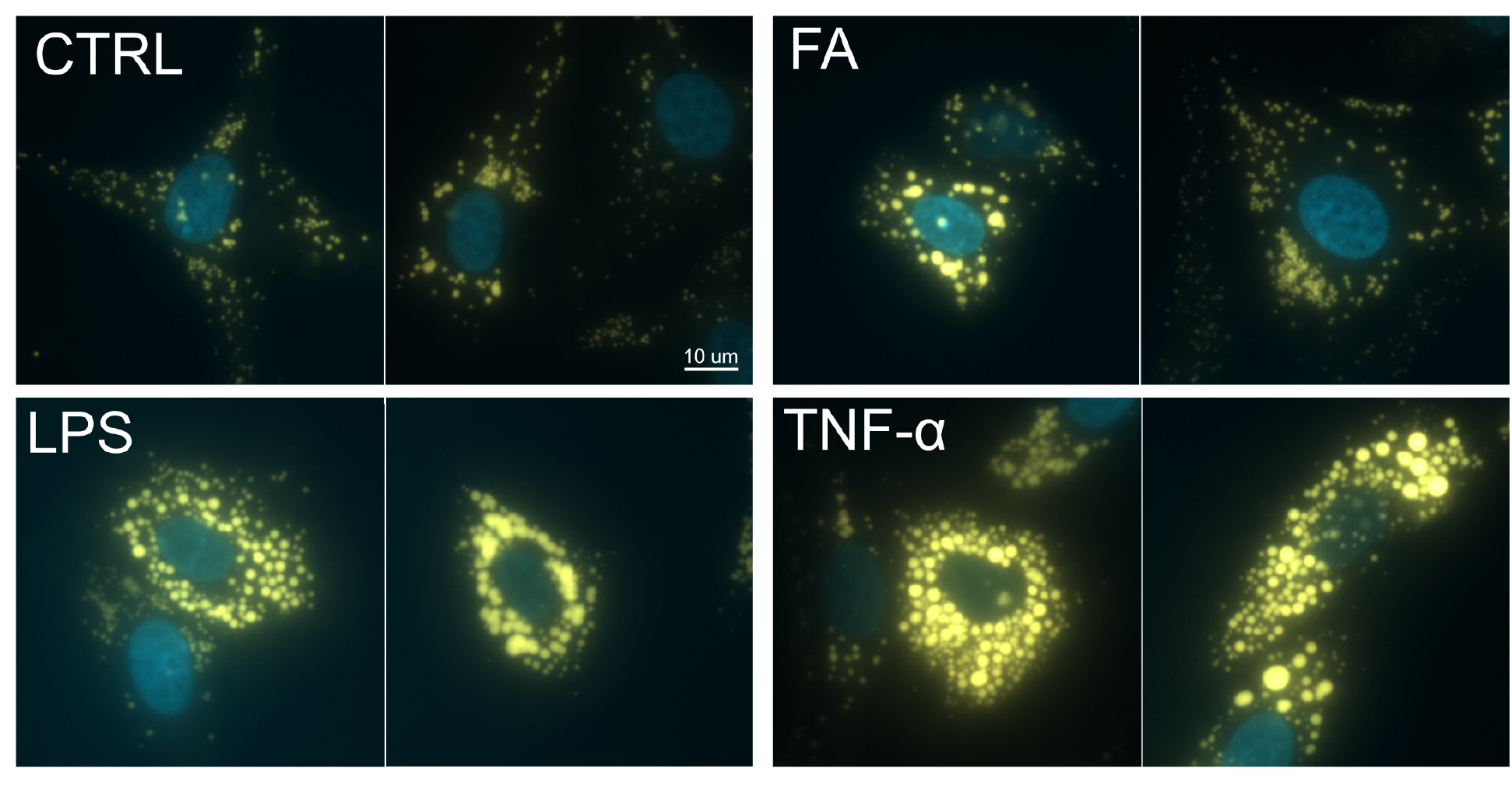
Lipid accumulation, also known as macro-vesicular steatosis, in dHepaRG hepatocytes both inherently as well as under stimulation with: oleic and palmitic fatty acids (FA), LPS in combination with the fatty acids, TNFα in combination with the fatty acids (yellow for LD540, blue for DAPI) to highlight the localization around nucleus. Each subplot shows two illustrative examples of cells.

**Supplementary Figure S4.**
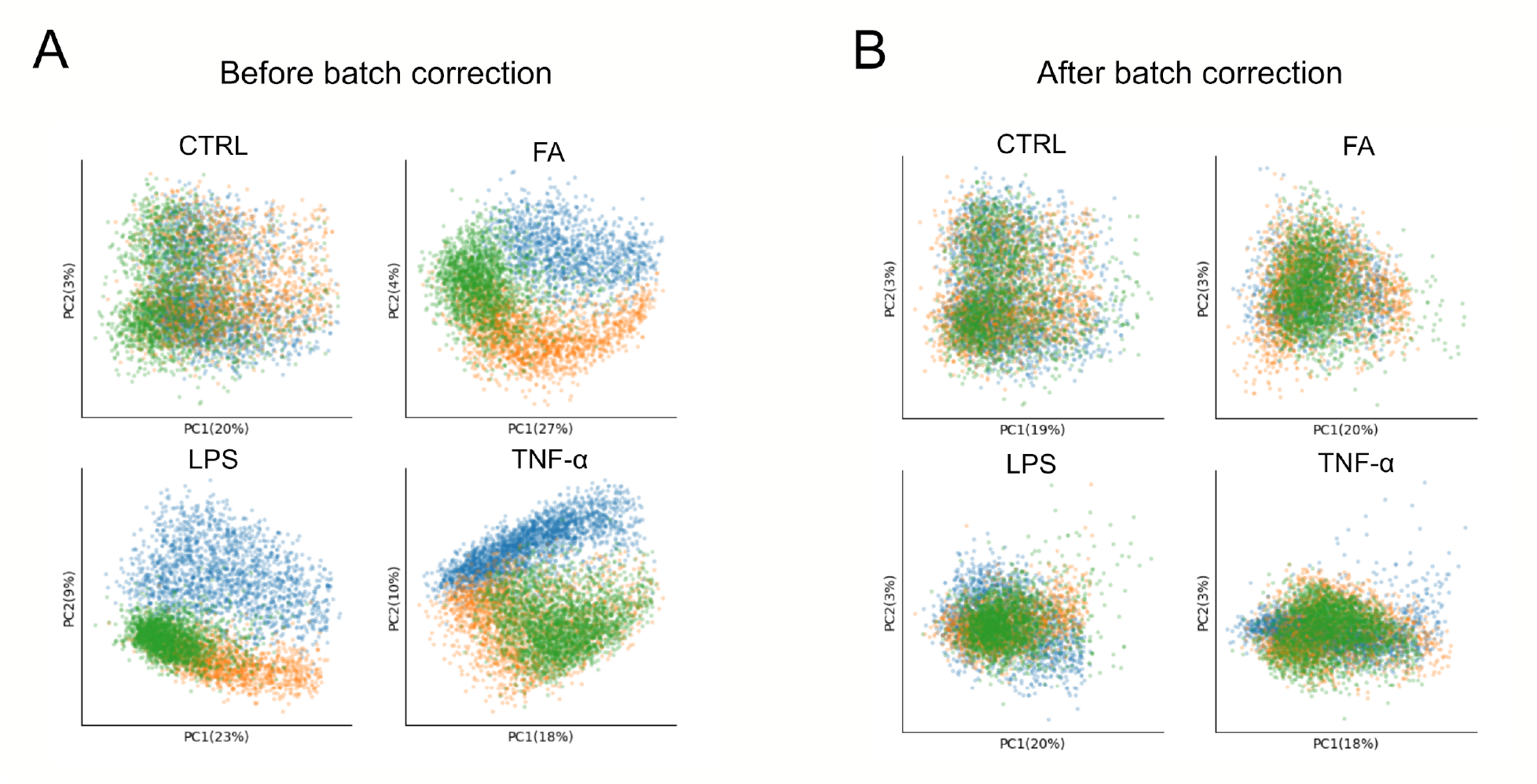
Batch correction of variability between technical replicates in the stimulated dHepaRG experiment by using the combat algorithm. The plots show a PCA plot of the single-cell metabolic profiles with cells color-coded according to the replicate. **A:** Before batch correction, **B:** after batch correction.

**SupplementaryFigure S5.**
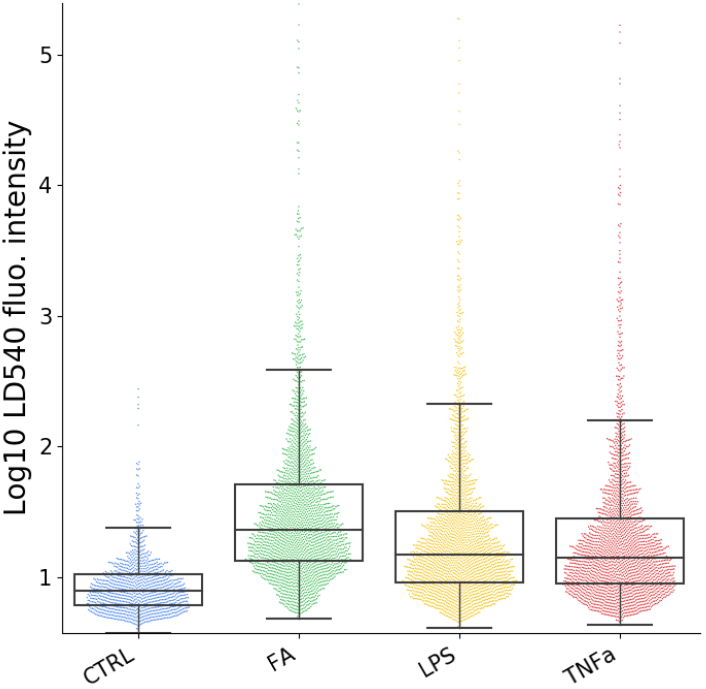
Swarming plots of single-cell LD540-fluorescent intensities of stimulated dHepaRG cells for each condition indicating increased lipid accumulation upon stimulation with FAs, LPS, and TNFα

**Supplementary Figure S6.**
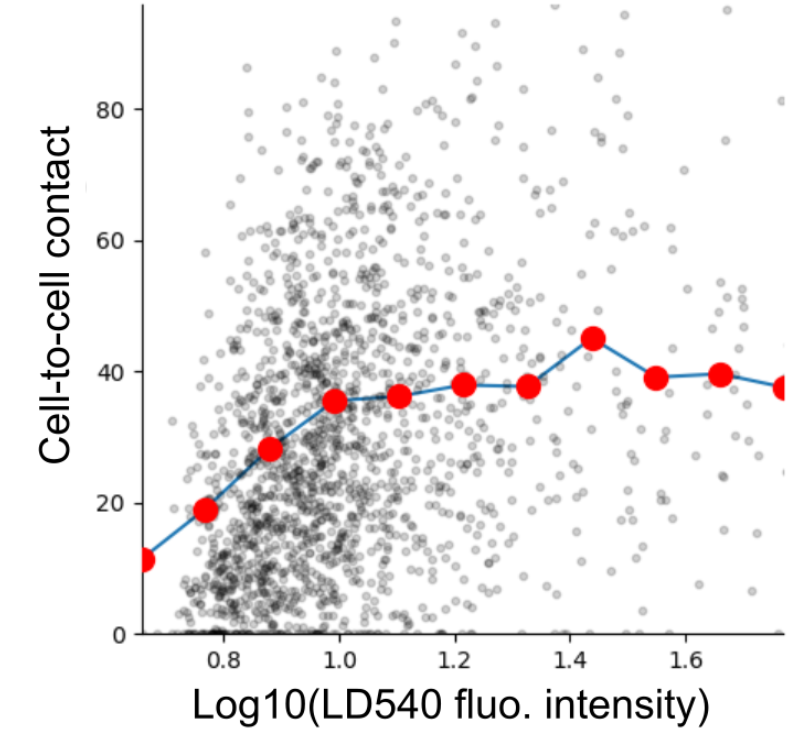
Scatter plot of the LD540 fluorescent intensity and cell-cell contact for one replicate of dHepaRG cells stimulated with TNFα in combination with oleic and linoleic acid (n=1830), with the Spearman r_s_ 0.34, p-value 9.18e-4. The LD540 intensities were thresholded at the 95% percentile. Red dots visualize average values for regularly-spaced bins of LD540 fluorescence intensity.

**Supplementary Figure S7.**
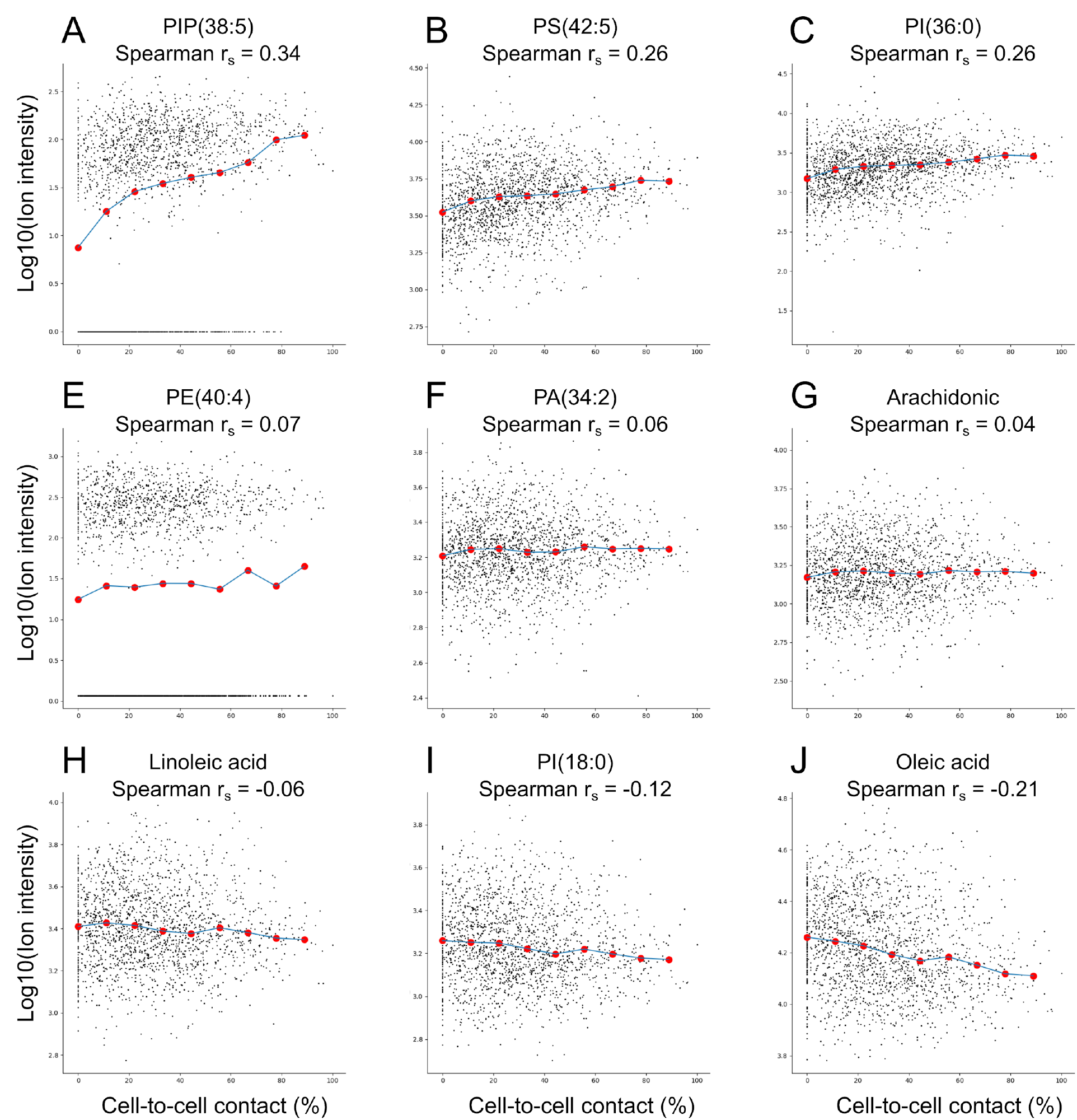
Observed relationship between local crowding and intracellular metabolite intensity. One dot represents one cell. Red dots represent average intensities for regular bins of local crowdedness. The data are coming from one replicate of the TNFα condition (1830 cells).

**Supplementary Figure S8.**
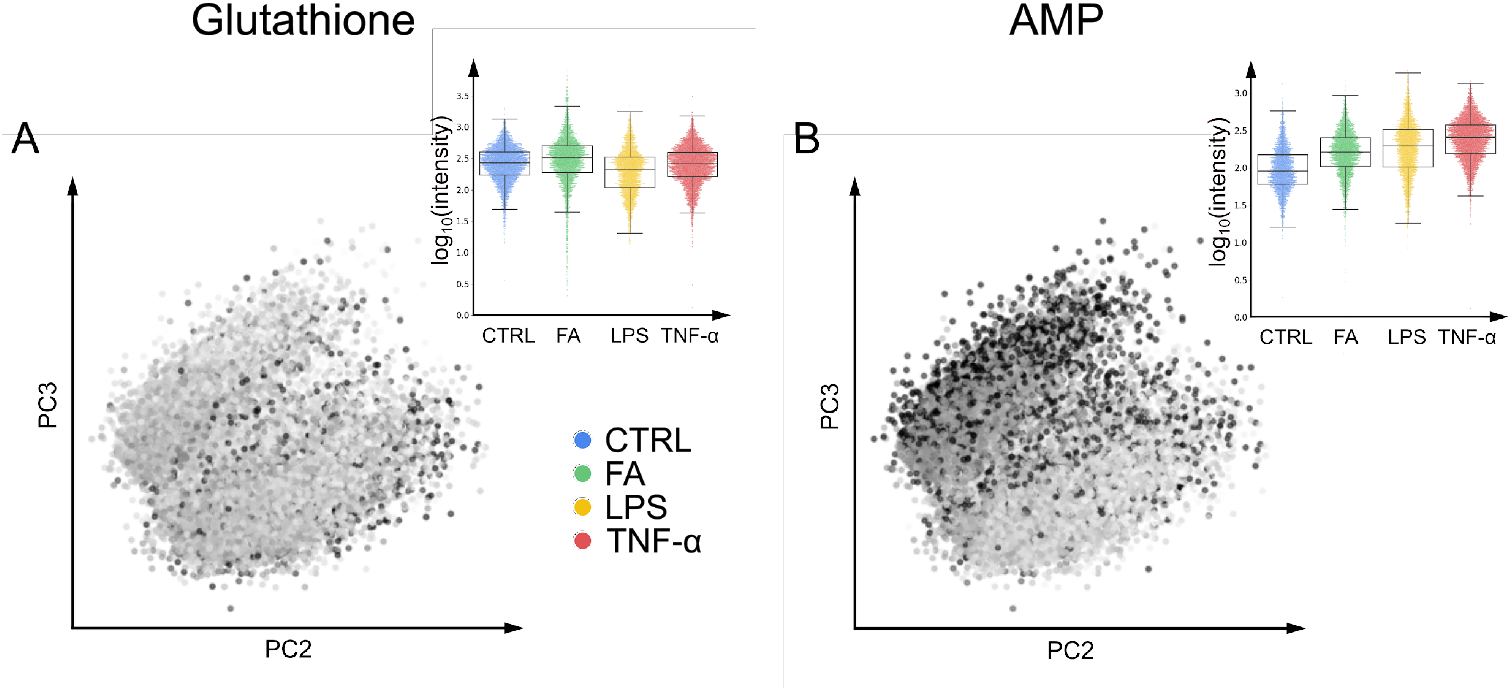
Mapping of intensities of the small molecules glutathione and AMP onto the principal components of z-scores of the single-cell metabolic profiles of dHepaRG cells from the CTRL, FA, LPS, and TNFα conditions (similar to Figure 3B-D).

**Supplementary Figure S9.**
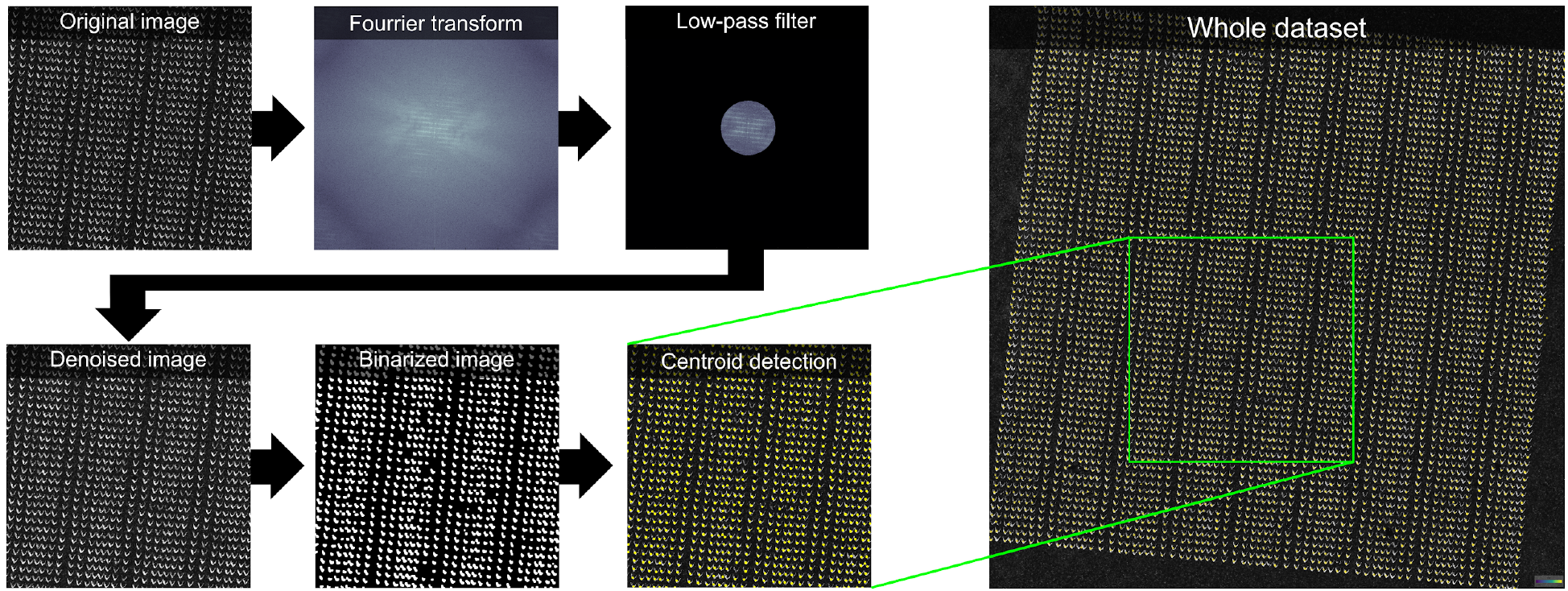
Illustration of the procedure for detection of laser ablation marks in post-MALDI microscopy images.

**Supplementary Figure S10.**
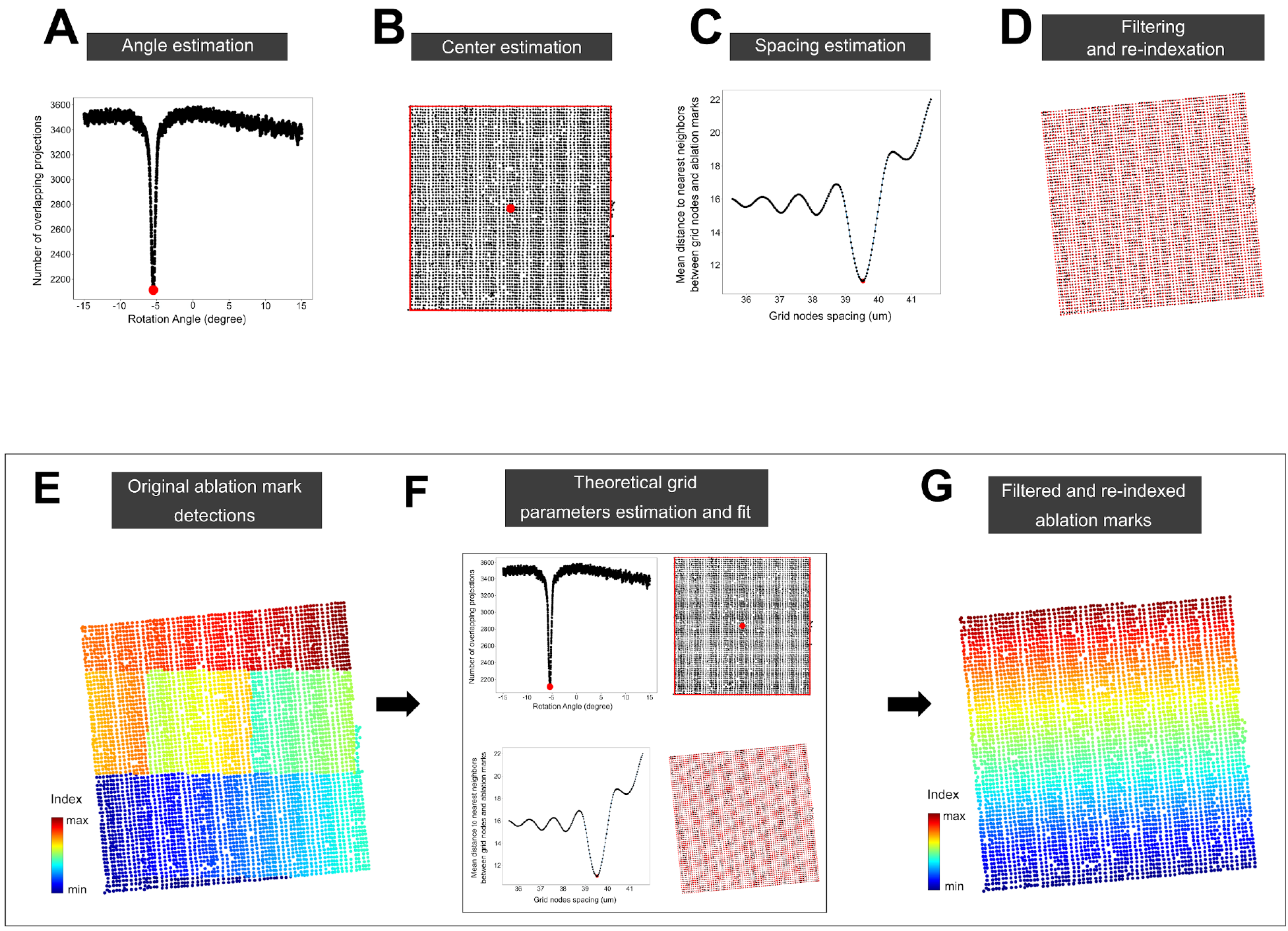
Illustration of the procedure for fitting a theoretical rectangular grid to the ablation marks segmented in the post-MALDI microscopy images. The three parameters of the grid are estimated using the ablation marks coordinates. In A, the angle is estimated by counting the number of non overlapping ablation marks coordinates projections of the X axis for different rotation angle. The minimum number of projection is reached for an alignment angle with the projection axis. In B, the center of the grid is estimated from the extrema of the ablation mark coordinates. The spacing of the grid nodes is estimated in C by measuring the mean distance to the nearest ablation mark to each grid node. The chosen grid node spacing is leads to smallest mean distance to the ablation mark coordinates. The re-indexing in D is done by choosing the closest ablation mark coordinates from the grid nodes constructed using the parameters defined before (the grid nodes are shown in red, their nearest ablation mark coordinates are shown in black). In E, the ablation mark coordinates are color coded by their index. An illustration of the different steps for fitting a grid onto the ablation mark coordinates as well as the re-indexing is shown in F. In G, the re-indexed ablation mark are shown. and re-indexing them to associate each detected ablation mark with a MALDI spectrum.

**Supplementary Figure S11.**
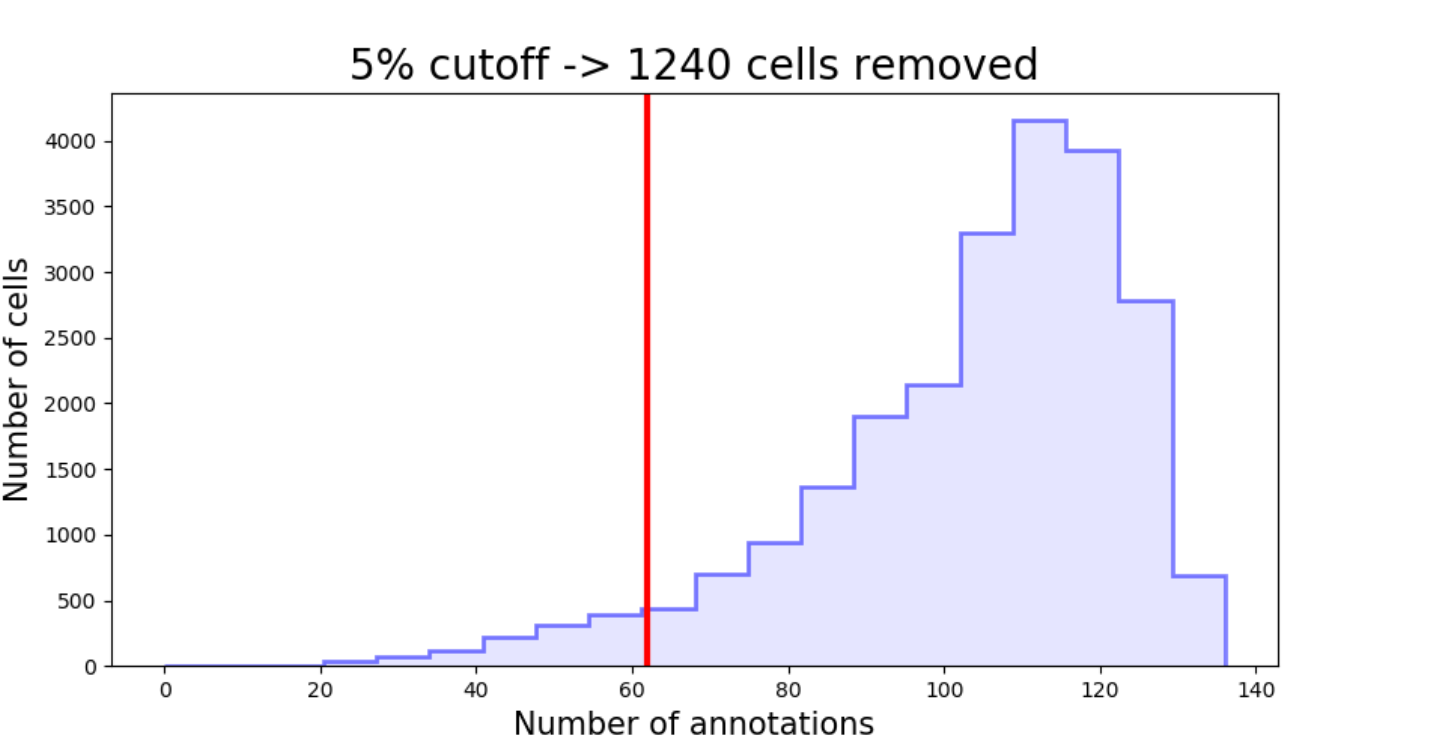
Filtering out poor quality cells for the dHepaRG experiment (Figure 4). The cells with low number of METASPACE annotations (less than the 5% lower percentile of the distribution) were removed.

## Supplementary Table

**Supplementary Table S1.** Experimental design and ambient conditions for matrix application and MALDI-imaging analysis of dHepaRG samples.

|  | Matrix application |  |  | MALDI-imaging |  |  |
| --- | --- | --- | --- | --- | --- | --- |
| Sample<br>("Condition" –<br>"Replicate<br>number") | Date; Time | Room<br>temperature<br>(°C) | Room<br>humidity<br>(%) | Date; Time | Room<br>temperature<br>(°C) | Room<br>humidity<br>(%) |
| FA_1 | 22.06.17; 1.40pm | 23.1 | 58 | 22.06.17; 2.01pm | 23.1 | 50 |
| LPS_1 | 22.06.17; 4.10pm | 23.2 | 60 | 22.06.17; 4.30pm | 23.7 | 49 |
| FA_2 | 24.06.17; 12.40am | 21.9 | 52 | 24.06.17; 1.00pm | 23.3 | 47 |
| LPS_2 | 24.06.17; 3.50pm | 21.9 | 53 | 24.06.17; 4.10pm | 23.5 | 47 |
| Untreated_1 | 25.06.17; 6.30am | 21.9 | 52 | 25.06.17; 7.00am | 23 | 48 |
| TNFa_1 | 25.06.17; 12.15am | 22.1 | 60 | 25.06.17; 3.30pm | 23.6 | 51 |
| FA_3 | 25.06.17; 6.30am | 21.9 | 58 | 25.06.17; 10.00am | 23.1 | 47 |
| TNFa_2 | 25.06.17; 9.15am | 21.9 | 60 | 25.06.17; 9.35am | 23.1 | 52 |
| Untreated_2 | 25.06.17; 12.41pm | 21.5 | 55 | 25.06.17; 4.00pm | 23.1 | 47 |
| Untreated_3 | 25.06.17; 12.41pm | 21.5 | 55 | 25.06.17; 1.00pm | 23.1 | 46 |
| LPS_3 | 28.06.17; 2.59pm | 21.1 | 63 | 28.06.17; 6.30pm | 23.3 | 48 |
| TNFa_3 | 30.06.17; 2.30pm | 21.1 | 50 | 30.06.17; 6.41pm | 22.6 | 44 |

## Supplementary Data (attached 920 as a separate file)

### Supplementary Data S1. LC-MS/MS validation of METASPACE metabolite annotations

Summary and detailed information about LC-MS/MS validation of METASPACE annotations including MS/MS and chromatographic information.

## References

Altschuler, S.J., and Wu, L.F. (2010). Cellular heterogeneity: do differences make a difference? Cell 141, 559–563.

Baker, T.C., Han, J., and Borchers, C.H. (2017). Recent advancements in matrix-assisted laser desorption/ionization mass spectrometry imaging. Curr. Opin. Biotechnol. 43, 62–69.

Beutler, B. (2004). Inferences, questions and possibilities in Toll-like receptor signalling. Nature 430, 257–263.

Bray, M.-A., Singh, S., Han, H., Davis, C.T., Borgeson, B., Hartland, C., Kost-Alimova, M., Gustafsdottir, S.M., Gibson, C.C., and Carpenter, A.E. (2016). Cell Painting, a high-content image-based assay for morphological profiling using multiplexed fluorescent dyes. Nat. Protoc. 11, 1757–1774.

Buck, M.D., Sowell, R.T., Kaech, S.M., and Pearce, E.L. (2017). Metabolic Instruction of Immunity. Cell 169, 570–586.

Carpenter, A.E., Jones, T.R., Lamprecht, M.R., Clarke, C., Kang, I.H., Friman, O., Guertin, D.A., Chang, J.H., Lindquist, R.A., Moffat, J., et al. (2006). CellProfiler: image analysis software for identifying and quantifying cell phenotypes. Genome Biol. 7, R100.

Do, T.D., Comi, T.J., Dunham, S.J.B., Rubakhin, S.S., and Sweedler, J.V. (2017). Single Cell Profiling Using Ionic Liquid Matrix-Enhanced Secondary Ion Mass Spectrometry for Neuronal Cell Type Differentiation. Anal. Chem. 89, 3078–3086.

Folch, J., Lees, M., and Sloane Stanley, G.H. (1957). A simple method for the isolation and purification of total lipides from animal tissues. J. Biol. Chem. 226, 497–509.

Fortin, J.-P., Cullen, N., Sheline, Y.I., Taylor, W.D., Aselcioglu, I., Cook, P.A., Adams, P., Cooper, C., Fava, M., McGrath, P.J., et al. (2017). Harmonization of cortical thickness measurements across scanners and sites. Neuroimage 167, 104–120.

Gluchowski, N.L., Becuwe, M., Walther, T.C., and Farese, R.V., Jr (2017). Lipid droplets and liver disease: from basic biology to clinical implications. Nat. Rev. Gastroenterol. Hepatol. 14, 343–355.

Gripon, P., Rumin, S., Urban, S., Le Seyec, J., Glaise, D., Cannie, I., Guyomard, C., Lucas, J., Trepo, C., and Guguen-Guillouzo, C. (2002). Infection of a human hepatoma cell line by hepatitis B virus. Proc. Natl. Acad. Sci. U. S. A. 99, 15655–15660.

Grun, D., and van Oudenaarden, A. (2015). Design and Analysis of Single-Cell Sequencing Experiments. Cell 163, 799–810.

Guillaume-Gentil, O., Rey, T., Kiefer, P., Ibanez, A.J., Steinhoff, R., Bronnimann, R., Dorwling-Carter, L., Zambelli, T., Zenobi, R., and Vorholt, J.A. (2017). Single-Cell Mass Spectrometry of Metabolites Extracted from Live Cells by Fluidic Force Microscopy. Anal. Chem. 89, 5017–5023.

Ibanez, A.J., Fagerer, S.R., Schmidt, A.M., Urban, P.L., Jefimovs, K., Geiger, P., Dechant, R., Heinemann, M., and Zenobi, R. (2013). Mass spectrometry-based metabolomics of single yeast cells. Proc. Natl. Acad. Sci. U. S. A. 110, 8790–8794.

Ilicic, T., Kim, J.K., Kolodziejczyk, A.A., Bagger, F.O., McCarthy, D.J., Marioni, J.C., and Teichmann, S.A. (2016). Classification of low quality cells from single-cell RNA-seq data. Genome Biol. 17, 29.

Jung, U.J., and Choi, M.-S. (2014). Obesity and its metabolic complications: the role of adipokines and the relationship between obesity, inflammation, insulin resistance, dyslipidemia and nonalcoholic fatty liver disease. Int. J. Mol. Sci. 15, 6184–6223.

Lee, M.-C.W., Lopez-Diaz, F.J., Khan, S.Y., Tariq, M.A., Dayn, Y., Vaske, C.J., Radenbaugh, A.J., Kim, H.J., Emerson, B.M., and Pourmand, N. (2014). Single-cell analyses of transcriptional heterogeneity during drug tolerance transition in cancer cells by RNA sequencing. Proc. Natl. Acad. Sci. U. S. A. 111, E4726–E4735.

Marioni, J.C., and Arendt, D. (2017). How Single-Cell Genomics Is Changing Evolutionary and Developmental Biology. Annu. Rev. Cell Dev. Biol. 33, 537–553.

Merrill, C.B., Basit, A., Armirotti, A., Jia, Y., Gall, C.M., Lynch, G., and Piomelli, D. (2017). Patch clamp-assisted single neuron lipidomics. Sci. Rep. 7, 5318.

Murphy, M.P., and O’Neill, L.A.J. (2018). Krebs Cycle Reimagined: The Emerging Roles of Succinate and Itaconate as Signal Transducers. Cell 174, 780–784.

Otsu, N. (1979). A threshold selection method from gray-level histograms. IEEE Trans. Syst. Man Cybern. 9, 62–66.

Palmer, A., Trede, D., and Alexandrov, T. (2016). Where imaging mass spectrometry stands: here are the numbers. Metabolomics 12, 1–3.

Palmer, A., Phapale, P., Chernyavsky, I., Lavigne, R., Fay, D., Tarasov, A., Kovalev, V., Fuchser, J., Nikolenko, S., Pineau, C., et al. (2017). FDR-controlled metabolite annotation for high-resolution imaging mass spectrometry. Nat. Methods 14, 57–60.

Passarelli, M.K., Pirkl, A., Moellers, R., Grinfeld, D., Kollmer, F., Havelund, R., Newman, C.F., Marshall, P.S., Arlinghaus, H., Alexander, M.R., et al. (2017). The 3D OrbiSIMS-label-free metabolic imaging with subcellular lateral resolution and high mass-resolving power. Nat. Methods 14, 1175–1183.

Patel, A.P., Tirosh, I., Trombetta, J.J., Shalek, A.K., Gillespie, S.M., Wakimoto, H., Cahill, D.P., Nahed, B.V., Curry, W.T., Martuza, R.L., et al. (2014). Single-cell RNA-seq highlights intratumoral heterogeneity in primary glioblastoma. Science 344, 1396–1401.

Pavlova, N.N., and Thompson, C.B. (2016). The Emerging Hallmarks of Cancer Metabolism. Cell Metab. 23, 27–47.

Pelkmans, L. (2012). Cell Biology. Using cell-to-cell variability--a new era in molecular biology. Science 336, 425–426.

Preibisch, S., Saalfeld, S., and Tomancak, P. (2009). Globally optimal stitching of tiled 3D microscopic image acquisitions. Bioinformatics 25, 1463–1465.

Protsyuk, I., Melnik, A.V., Nothias, L.-F., Rappez, L., Phapale, P., Aksenov, A.A., Bouslimani, A., Ryazanov, S., Dorrestein, P.C., and Alexandrov, T. (2018). 3D molecular cartography using LC-MS facilitated by Optimus and ‘ili software. Nat. Protoc. 13, 134–154.

Ress, C., and Kaser, S. (2016). Mechanisms of intrahepatic triglyceride accumulation. World J. Gastroenterol. 22, 1664–1673.

Rubakhin, S.S., Lanni, E.J., and Sweedler, J.V. (2013). Progress toward single cell metabolomics. Curr. Opin. Biotechnol. 24, 95–104.

Russell, A.B., Trapnell, C., and Bloom, J.D. (2018). Extreme heterogeneity of influenza virus infection in single cells. Elife 7.

Sharma, U., and Rando, O.J. (2017). Metabolic Inputs into the Epigenome. Cell Metab. 25, 544–558.

Sharon, G., Garg, N., Debelius, J., Knight, R., Dorrestein, P.C., and Mazmanian, S.K. (2014). Specialized metabolites from the microbiome in health and disease. Cell Metab. 20, 719–730.

da Silva-Santi, L.G., Antunes, M.M., Caparroz-Assef, S.M., Carbonera, F., Masi, L.N., Curi, R., Visentainer, J.V., and Bazotte, R.B. (2016). Liver Fatty Acid Composition and Inflammation in Mice Fed with High-Carbohydrate Diet or High-Fat Diet. Nutrients 8.

Soltwisch, J., Kettling, H., Vens-Cappell, S., Wiegelmann, M., Muthing, J., and Dreisewerd, K. (2015). Mass spectrometry imaging with laser-induced postionization. Science 348, 211–215.

Spandl, J., White, D.J., Peychl, J., and Thiele, C. (2009). Live cell multicolor imaging of lipid droplets with a new dye, LD540. Traffic 10, 1579–1584.

Sumner, L.W., Amberg, A., Barrett, D., Beale, M.H., Beger, R., Daykin, C.A., Fan, T.W.-M., Fiehn, O., Goodacre, R., Griffin, J.L., et al. (2007). Proposed minimum reporting standards for chemical analysis Chemical Analysis Working Group (CAWG) Metabolomics Standards Initiative (MSI). Metabolomics 3, 211–221.

Wellen, K.E., and Thompson, C.B. (2012). A two-way street: reciprocal regulation of metabolism and signalling. Nat. Rev. Mol. Cell Biol. 13, 270–276.

Wishart, D.S., Knox, C., Guo, A.C., Eisner, R., Young, N., Gautam, B., Hau, D.D., Psychogios, N., Dong, E., Bouatra, S., et al. (2009). HMDB: a knowledgebase for the human metabolome. Nucleic Acids Res. 37, D603–D610.

Wolf, M.J., Adili, A., Piotrowitz, K., Abdullah, Z., Boege, Y., Stemmer, K., Ringelhan, M., Simonavicius, N., Egger, M., Wohlleber, D., et al. (2014). Metabolic activation of intrahepatic CD8+ T cells and NKT cells causes nonalcoholic steatohepatitis and liver cancer via cross-talk with hepatocytes. Cancer Cell 26, 549–564.

Zenobi, R. (2013). Single-cell metabolomics: analytical and biological perspectives. Science 342, 1243259.

