## Supplementary Data S1 for "Spatial single-cell profiling of intracellular metabolomes *in situ*"

### LC-MS/MS validation of METASPACE metabolite annotations

### Summary of the LC-MS/MS validation

| Molecular formula | Cell type | METASPACE annotation | Validated using LC-MS/MS as | Validation method | Summary of the MS/MS validation |
|---|---|---|---|---|---|
| C18H32O2 | dHepaRG | Linoleic acid* | Linoleic acid | Lipidomics (negative mode) | **Standard:** 9-cis,12-cis-Linoleic acid, Sigma L1376 [HMDB0001388] |
| C18H34O2 | dHepaRG | Oleic acid* | Oleic acid | Lipidomics (negative mode) | **Standard:** cis-9-Octadecenoic acid, Sigma O1008 [HMDB0000207] |
| C10H14N5O7P | dHepaRG | AMP* | AMP | Metabolomics (negative mode) | **SL:** EMBL-MCF spectral library |
| C47H82O16P2 | dHepaRG | PIP(38:5) | PIP(20:4_18:1); PIP(22:5_16:0) | Lipidomics (negative mode) | **HD:** PI<br>**SD:** 20:4,18:1;22:5,16:0 FA |
| C53H100O6 | dHepaRG | TG(50:1) | TG(16:0_18:1_16:0) | Lipidomics (positive mode) | **SL:** EMBL-MCF spectral library<br>**SD:** 16:0, 18:1 FA |
| C55H102O6 | dHepaRG | TG(52:2) | TG(16:0_18:1_18:1) | Lipidomics (positive mode) | **SL:** EMBL-MCF spectral library<br>**SD:** 16:0, 18:1 FA |
| C43H81O13P | HeLa and NIH3T3 | PI(34:1) | PI(16:0_18:1) | Lipidomics (negative mode) | **SD:** 16:0, 18:1 FA |
| C45H78NO8P | HeLa and NIH3T3 | PE(40:6)* | PE(18:0_22:6) | Lipidomics (positive mode) →<br>Lipidomics (negative mode) → | **HD:** PE<br>**SD:** 18:0, 22:6 FA |
| C10H17N3O6S | dHepaRG | Glutathione | Glutathione | Metabolomics (negative mode) | **SL:** EMBL-MCF spectral library |

*** indicates ambiguity due to the existence of structural isomers**

**SL: MS/MS spectral library**
**HD: Head group MS/MS**
**SD: Fatty acid (FA) side chain**

**METASPACE annotation: C18H32O2, -H, m/z 279.232, Linoleic acid***

**Standard used** : **9-cis,12-cis-Linoleic acid,** Sigma **L1376 [HMDB0001388]**

**Validated as** : **Linoleic acid** using LC-MS/MS method **Lipidomics** (negative mode)

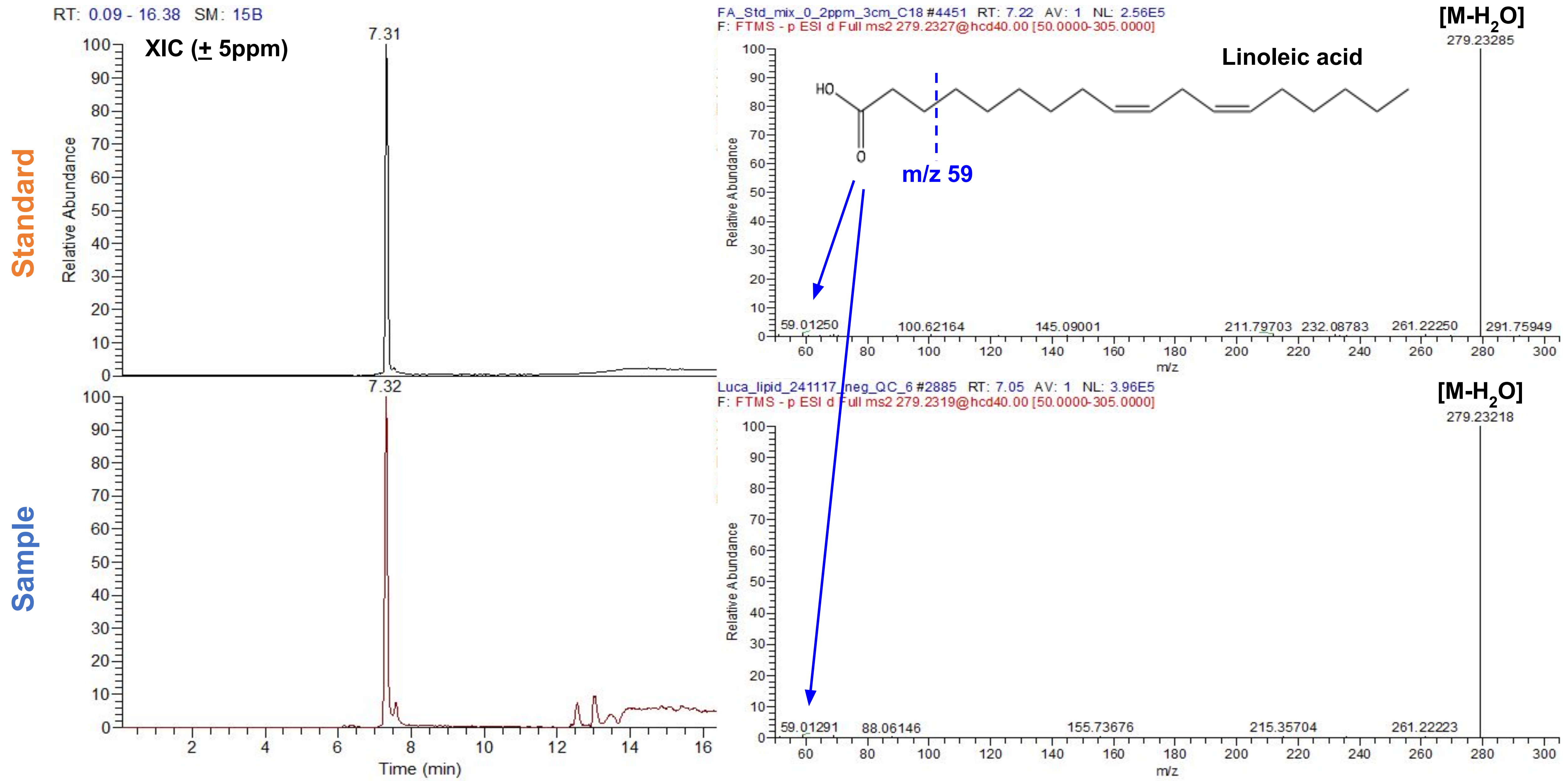

**METASPACE annotation: C18H34O2, -H, m/z 281.2558, Oleic acid***

**Standard used** : **cis-9-Octadecenoic acid,** Sigma O1008 [HMDB0000207]

**Validated as** : **Oleic acid** using LC-MS/MS method **Lipidomics** (negative mode)

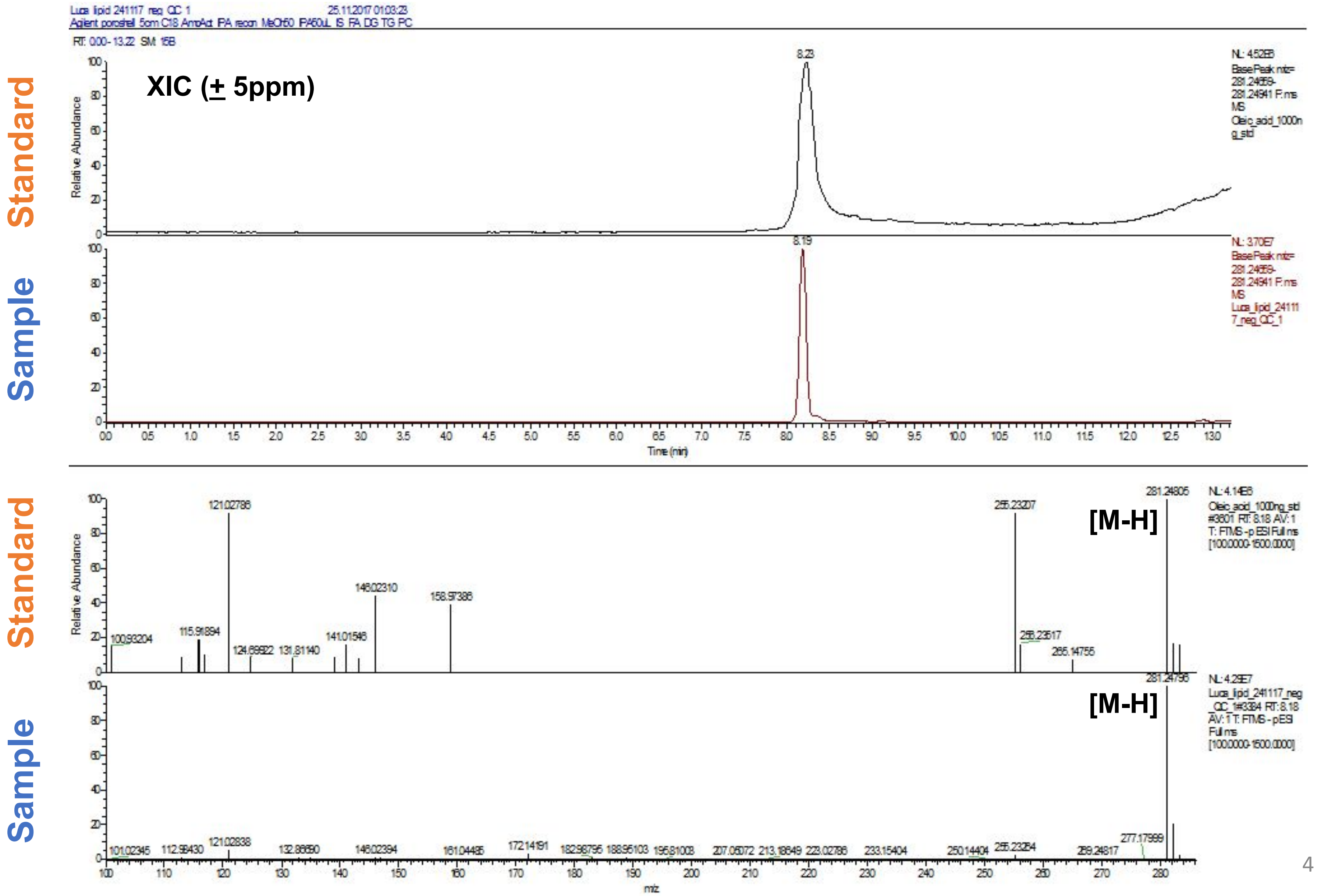

**METASPACE annotation: C10H14N5O7P, -H, m/z 346.0569, AMP***

**Validated as** : **AMP** using LC-MS/MS method **Metabolomics** (negative mode)

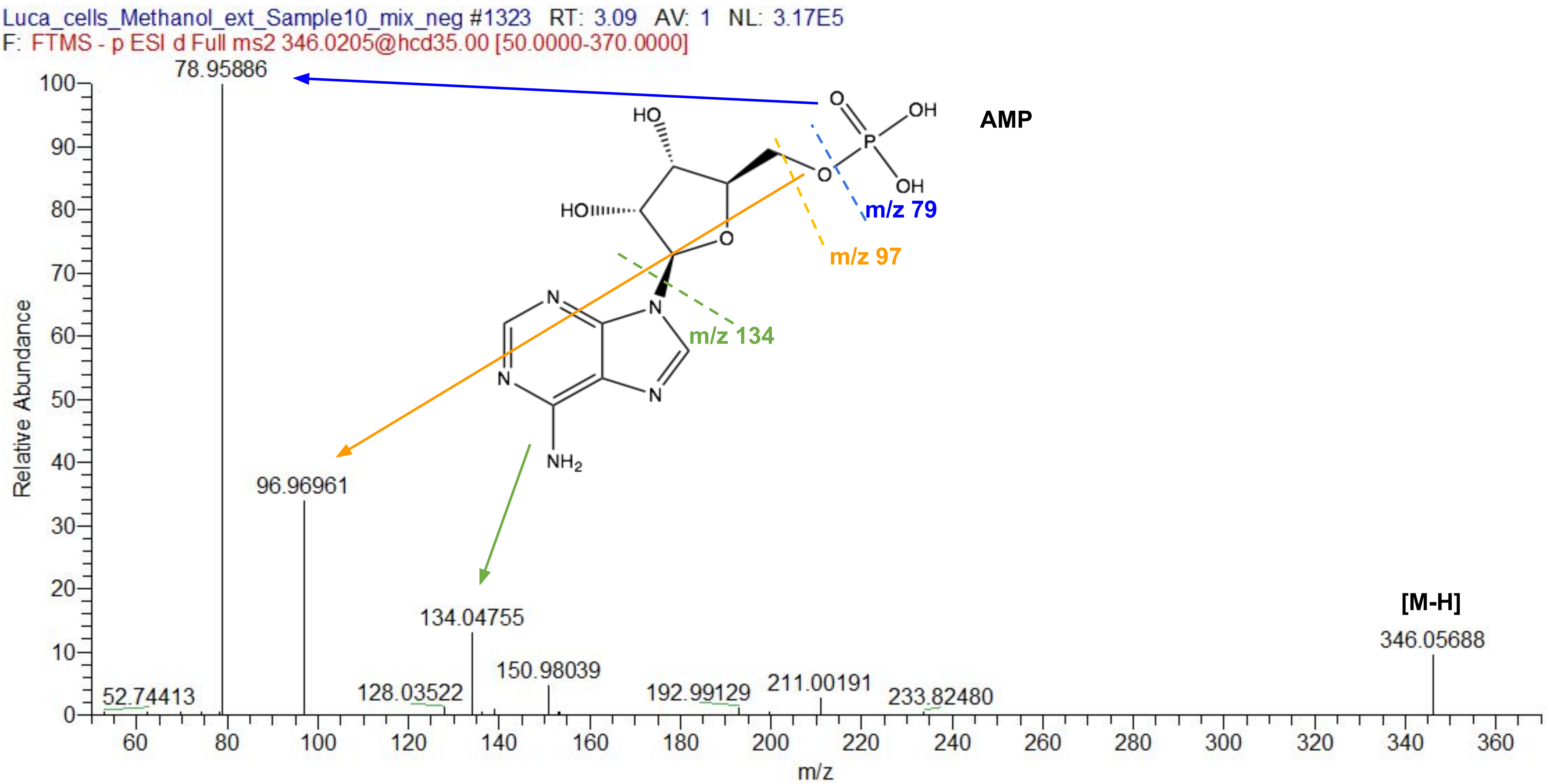

**METASPACE annotation: C47H82O16P2, -H , m/z 963.512, PIP(38:5)**

**Validated as : PIP(20:4_18:1) and PIP(22:5_16:0)** using LC-MS/MS method **Lipidomics** (negative mode)

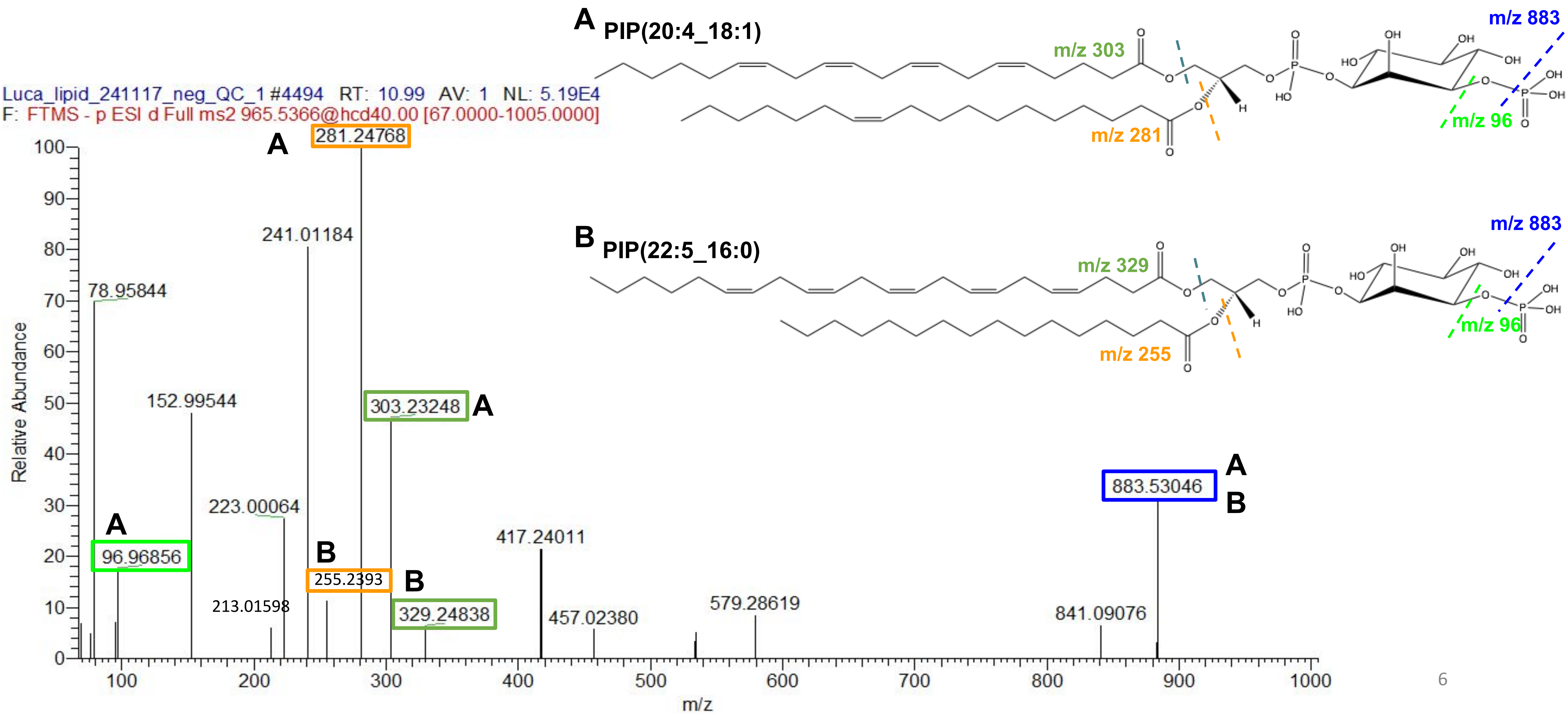

**METASPACE annotation: C53H100O6, +H, m/z 833.7520, TG(50:1)**

**Database used** : **EMBL-MCF** spectral library and LipidBlast search (in Progenesis software)

**Validated as** : **TG(16:0_18:1_16:0)** using LC-MS/MS method **Lipidomics** (positive mode)

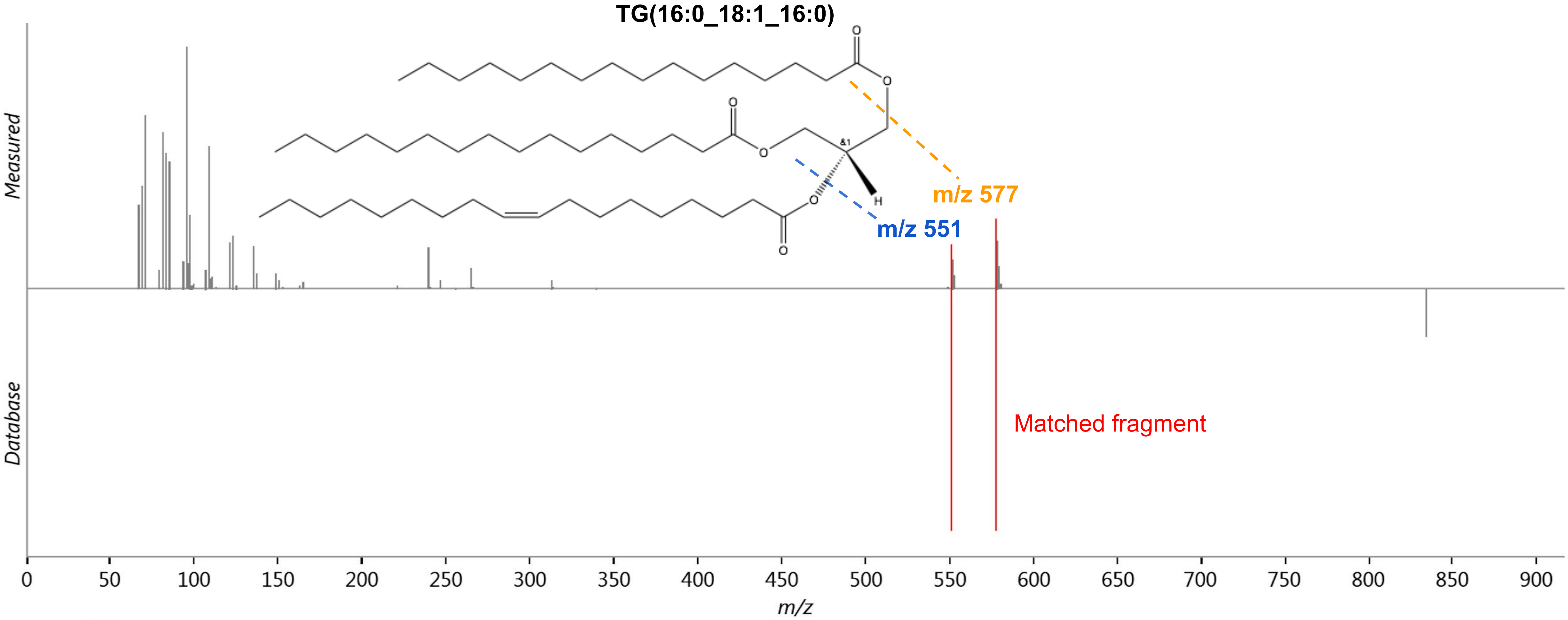

**METASPACE annotation: C55H102O6, +H, m/z 859.7676, TG(52:2)**

**Database used** : **EMBL-MCF** spectral library and LipidBlast search (in Progenesis software)

**Validated as** : **TG(16:0_18:1_18:1)** using LC-MS/MS method **Lipidomics** (positive mode)

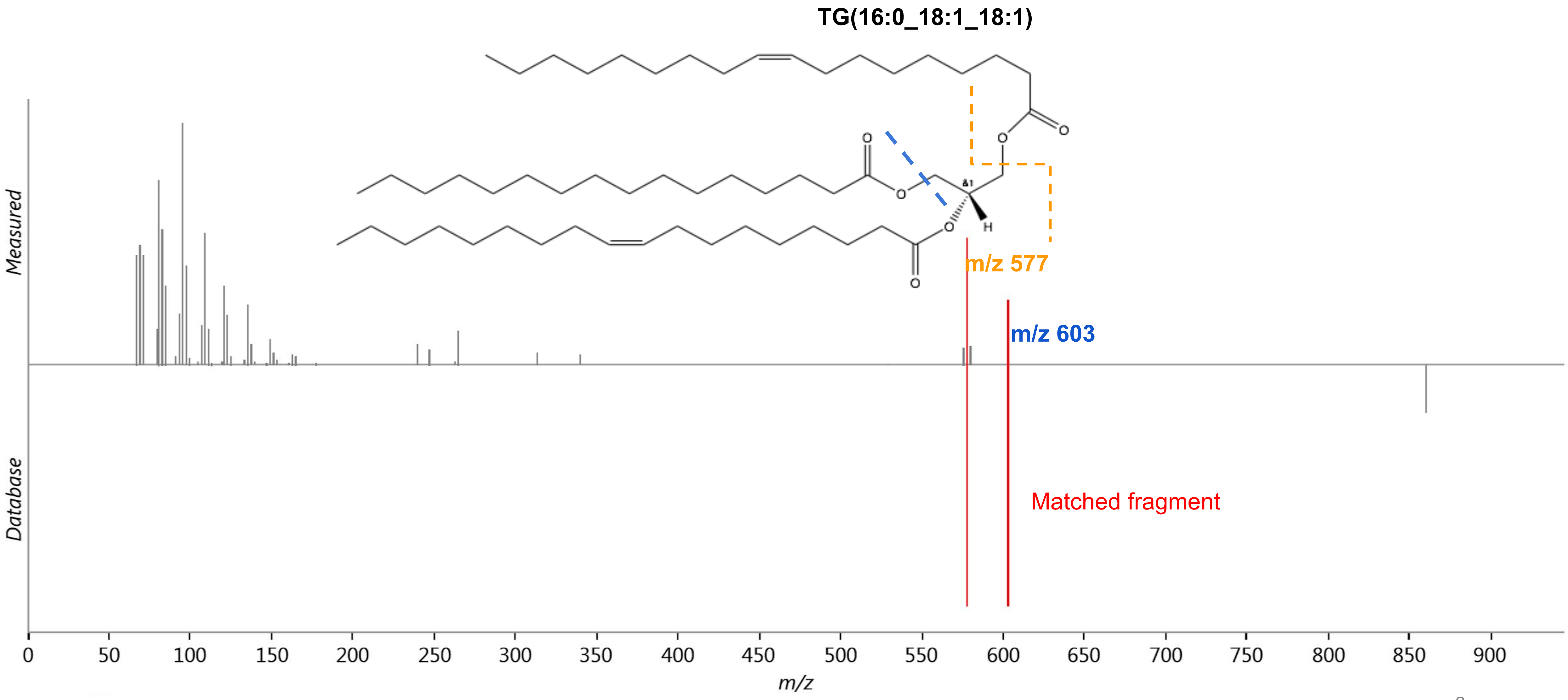

**METASPACE annotation: C43H81O13P, -H, m/z 835.53365, PI(34:1)**

**Validated as : PI(16:0_ 18:1)** using LC-MS/MS method **Lipidomics** (positive mode)

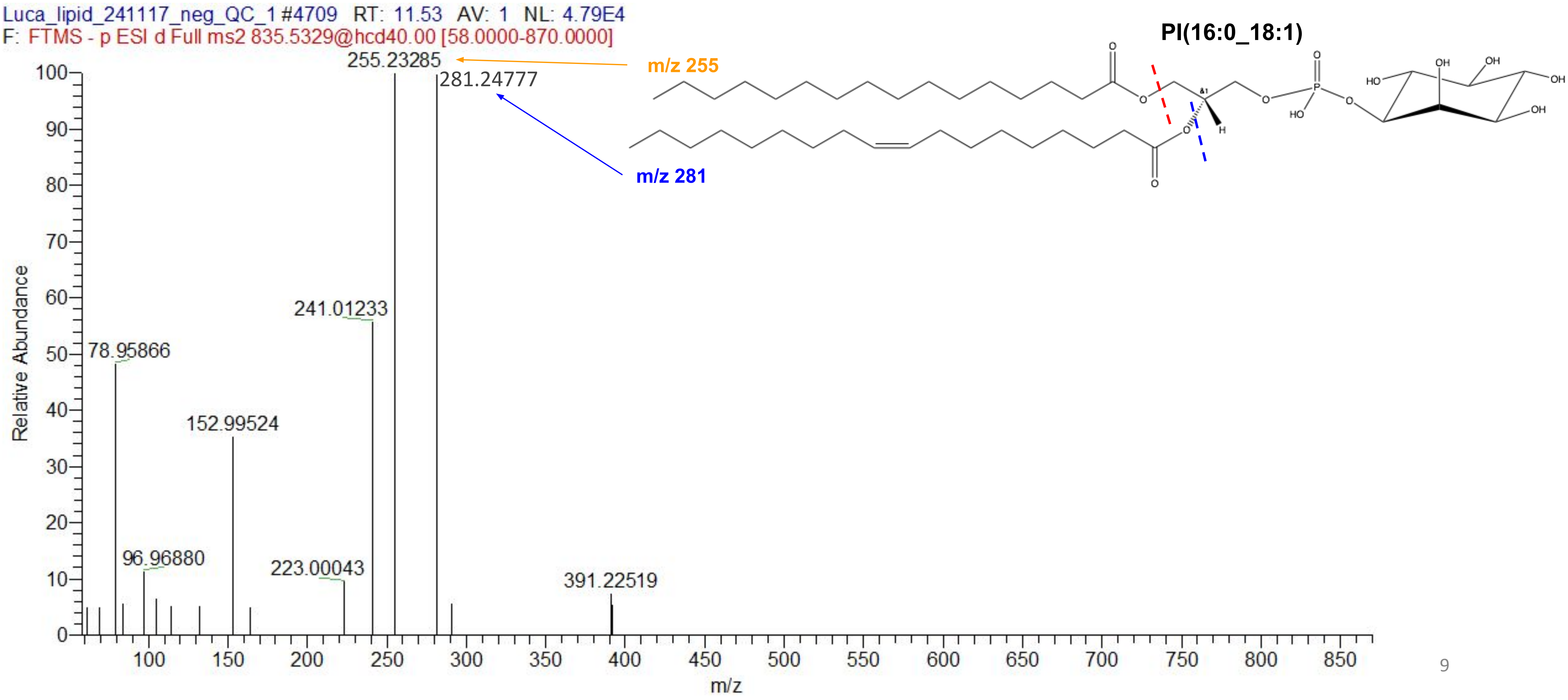

**METASPACE annotation: C45H78NO8P, +H, m/z 792.55433 PE(40:6)***

**Validated as** : **PE(40:6)** using LC-MS/MS method **Lipidomics** (positive mode)

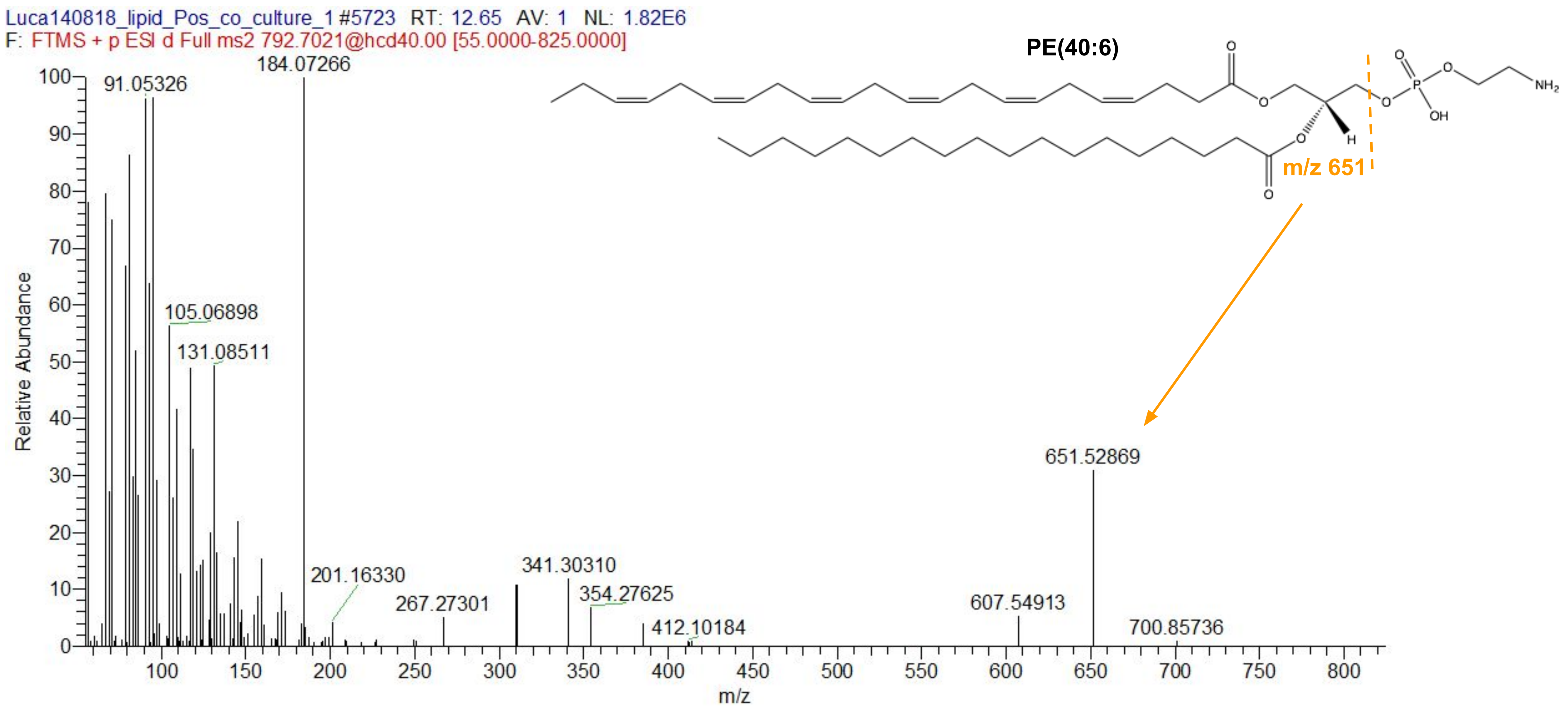

**METASPACE annotation: C45H78NO8P, -H, m/z 790.5386, PE(40:6)***

**Validated as : PE(18:0_22:6)** using LC-MS/MS method **Lipidomics** (negative mode)

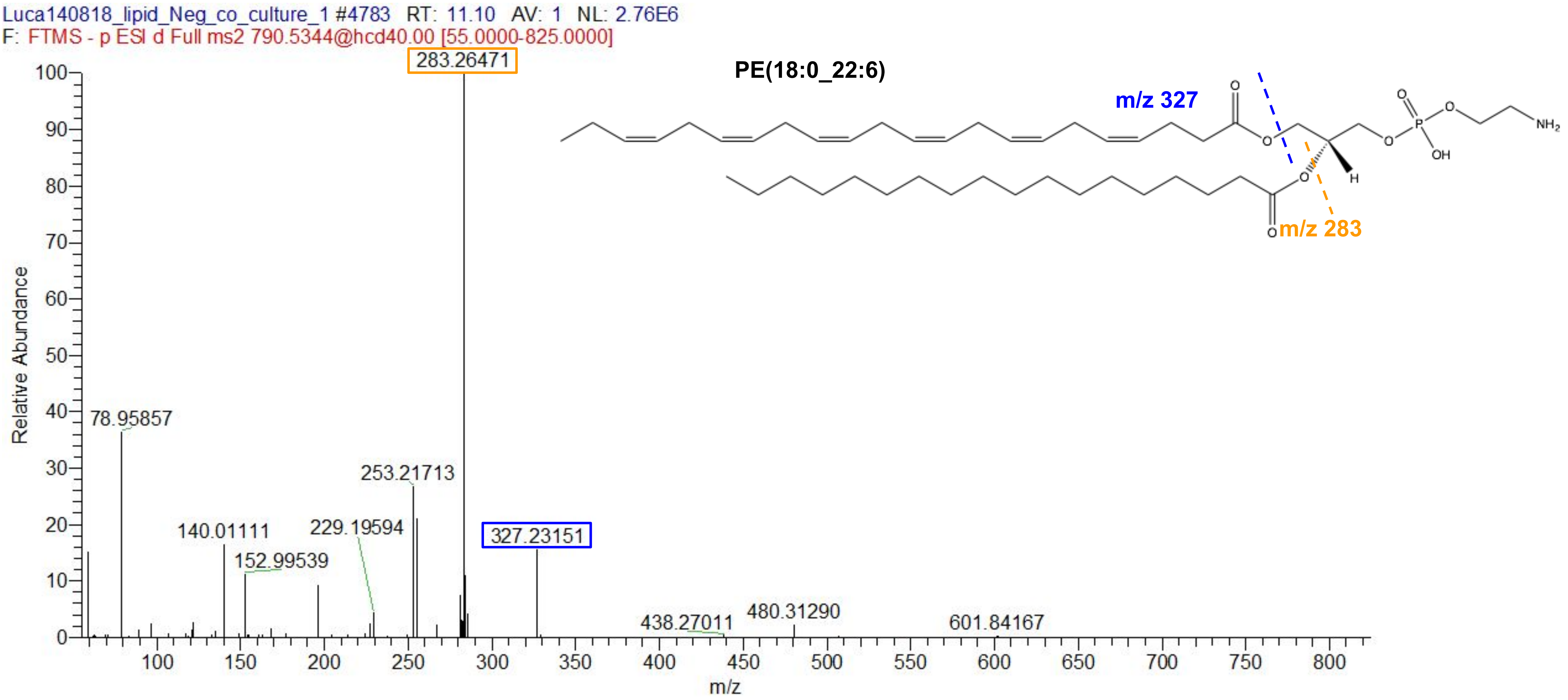

**METASPACE annotation: C10H17N3O6S, -H, m/z 306.0764, Glutathione**

**Validated as** : **Glutathione** using LC-MS/MS method **Metabolomics** (negative mode)

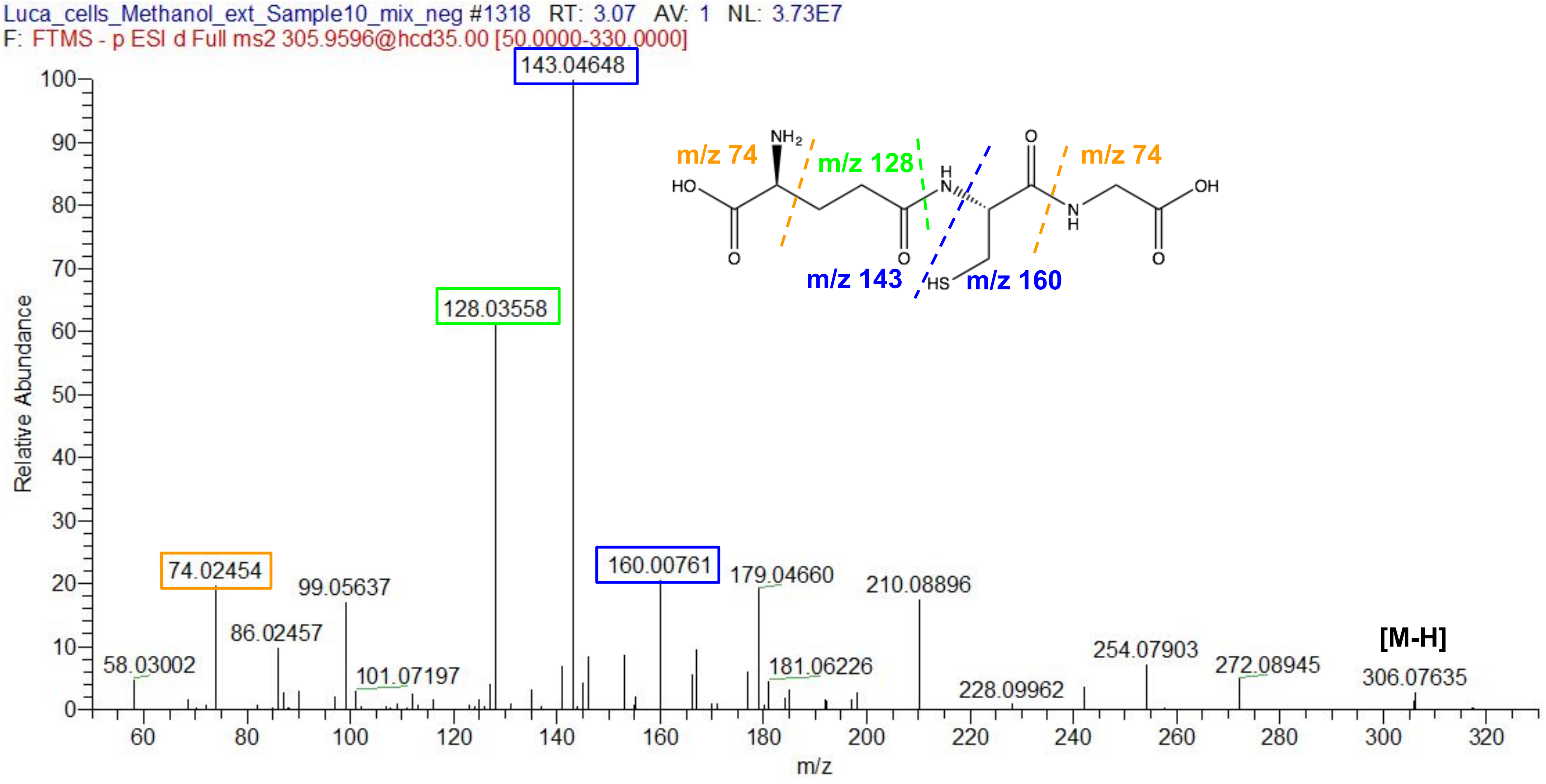
